# The unique structure and replication mode of the replication system of *Klebsiella pneumoniae* plasmid pIGMS31

**DOI:** 10.1101/2022.12.05.519187

**Authors:** Paweł Wawrzyniak, Sylwia Barańska, Pablo Hernández, Maciej Dylewski, Violetta Cecuda-Adamczewska, Łukasz Stypiński, Paulina Kabaj, Agata Gościk, Dariusz Bartosik

## Abstract

Due to their relatively small size and the lack of housekeeping genes, bacterial plasmids are very convenient models for studying DNA replication. So, for a long time, they had been intensively studied in this regard. Unfortunately, only a limited number of model plasmid replication systems were analyzed in detail. In the era of high-throughput DNA sequencing, we are faced with an increasing gap in our knowledge of bacterial plasmid replication and a rapidly growing number of deposited new plasmid genome sequences. For this reason, we decided to investigate the replication system of the pIGMS31 plasmid, a representative of the newly described pHW126-like plasmids family. They are small replicons isolated from different clinical and environmental strains of Gamma proteobacteria. Whole shares unique replication modules with no significant similarities to known model plasmid replication systems. In this study, we identified and characterized the basic elements of the pIGMS31 replication module. Studies on regulatory mechanisms of replication initiation of this plasmid as well as on pIGMS31 replication mode were also performed. We revealed that the pIGMS31 replication module is composed of elements typical for both *theta* and rolling circle replicons. This mosaic structure is reflected in the unique course of replication of this plasmid, with both modes of replication. What is more, our results led us to conclude that pHW126-like plasmids, despite DNA sequence similarity, are a highly diverse group of replicons.

## Introduction

Bacterial plasmids are a specific group of mobile genetic elements (MGEs). They exist as autonomous circular (less often linear) double-stranded DNA particles (Funnel and Phillips, 2004). Extensive research on plasmid biology follow from their role as genetic carriers transferring antibiotic resistance and virulence genes between clinical and environmental strains of different bacterial species (Johnson et al., 2012; Adamczuk et al., 2015; Pedersen et al., 2020; Dimitriu, 2022). Due to their relatively small size and the lack of housekeeping genes (they are not required by their hosts), bacterial plasmids are convenient research models that have been highly valuable in uncovering the molecular basis of many cellular processes, including DNA replication (Thomas et al., 1998; del Solar et al., 1998; Rajewska et al., 2012). Bacterial plasmids are also widely used in molecular biology, biotechnology, veterinary, and medicine. They are applied to the construction of different genetic tools, from DNA cloning vectors and heterologous gene overexpression vectors to antigen or therapeutic gene carriers used as DNA vaccines or gene therapy vehicles respectively (Camps, 2010; Faurez et al., 2010; Zaleski et al., 2017).

Regardless of their high genetic load diversity, the whole of bacterial plasmids has a set of different-type genetic modules indispensable for their maintenance (Thomas et al., 1998). The basic element of this genetic core is the REP module, responsible for the initiation of plasmid DNA vegetative replication (del Solar et al., 1998). Each of the REP modules is composed of *ori* (*origin*), where DNA replication starts, and initiation factor, RNA or Rep(s) protein(s) (del Solar et al., 1998). In general, based on the type of replication mechanism, circular bacterial plasmids can be divided into three groups: *theta, sigma* (rolling circle, RC) (so called because of the characteristic shape of their replication intermediates, RIs, resembling Greek letters *θ* or *δ* respectively), or strand displacement (D-loop) plasmids (del Solar et al., 1998). In fact, very few plasmid REP systems have been subjected to more detailed molecular analysis (Lilly and Camps, 2015, Ruiz-Masó et al., 2015). What is more, the rapid increase of DNA sequencing data results in a great variety of new plasmid genomes deposited in databases. Some of them encode Rep proteins with only weak or no sequence similarity to known replication initiators. It makes them very interesting research models for studying the heterogenicity of bacterial plasmid replicon systems. Unfortunately, we are not able to experimentally analyze each of the newly sequenced plasmids REP. However, it is definitely worth paying attention to those with high novelty, as well as those with application potential.

Recently, a new pHW126-like plasmid family was distinguished (Rozhon et al., 2010). They are small narrow-host-range (NHR) cryptic replicons from Gamma proteobacteria. They are composed of only two genetic modules, related REP modules, and different types of MOB modules responsible for mobilization for conjugal transfer. Some initial experiment results suggest that pHW126, an archetype of these plasmids, replicates via RC mode (Rozhon et al., 2010; Rozhon et al. 2011; Rozhon et al. 2012). But the Rep protein from pHW126 has no significant amino acid sequence similarity with known RC initiators nor with any of the other well-defined plasmid Rep protein types (Rozhon et al., 2010).

In the Institute of Biotechnology and Antibiotics (Warsaw, Poland), two other representatives of the pHW126-like plasmid family were isolated from *Klebsiella pneumoniae* clinical strain: pIGRK and pIGMS31 (Smorawińska et al., 2012). Both plasmids are deprived of antibiotic resistance or virulence genes, but we demonstrated that they can act as mobile carriers and transfer genetic material even to distantly related bacterial species in the conjugal transfer process (Smorawińska et al., 2012). What is more, REP modules of pIGRK and pIGMS31 or their elements were used as components of bacterial expression vectors. For example, the pIGRK *rep* gene promoter, cloned in the pIGDMKUH vector, was used to control the expression of human growth hormone (hGH) in industrially used *Escherichia coli* host strain (Płucienniczak et al., 2005; Płucienniczak et al., 2011; Mikiewicz et al., 2019).

*In silico* predictions revealed another interesting perspective on the use of REP modules from pIGRK and pIGMS31. The extremally low GC content of their DNA results in a minimal number of immunostimulatory sequences (ISS) (Van Uden and Raz, 2000). This suggests that pIGRK and pIGMS31 REP modules can be used as vectors for vaccine or gene therapy (Zaleski et al., 2017). We assume that they could escape from degradation and could provide sufficient heterologous gene expression in target cells. These predictions were proved by our preliminary experiments (for more detail see Supplementary Figure S1). Such advanced use requires meticulous characterization of plasmid REP modules to avoid genetic construct instability, undesirable DNA recombination events, or toxic effects, and to predict the optimal localization of therapeutic gene expression cassette insertion site (Faurez et al., 2010; Zaleski et al., 2017).

Tacking together, the unique character of pIGRK and pIGMS31 REP modules, distinct from known plasmid replication systems, and high application potential we decided to perform their molecular and functional analyses. In a last report, replication module of pIGRK plasmid was characterized in detail (Wawrzyniak et al., 2019; Nowak et al., 2021). Here we described a comprehensive analysis of pIGMS31 REP structural elements and its regulatory mechanisms. What is more, for the first time we investigated the replication mode of pHW126-like plasmid by direct visualization of DNA replication intermediates (RIs) arising during plasmid pIGMS31 genome replication. As a result, we revealed that the mosaic structure of the pIGMS31 REP (composed of RC-like and *theta*-like elements) is reflected in the unique course of replication of this plasmid, most probably, with both modes of replication. In addition, the presence of significant differences between known pHW126-like replicons was described.

## Materials and Methods

### Bacterial strains, plasmids, and culture conditions

Bacterial strains and plasmids used in this study are listed in Supplementary Table S1 All strains were cultured at 37°C in lysogeny broth (LB) medium (tryptone 10.0 g/l, yeast extract 5.0 g/l, and NaCl 5.0 g/l; pH 7.2–7.5). When necessary, the medium was supplemented with appropriate antibiotics at the following concentrations: ampicillin (Ap) – 100 μg/ml, kanamycin (Km) – 25 μg/ml [for *E. coli* BL21(DE3)] or 50 μg/ml (for other strains), or chloramphenicol (Cm) – 34 μg/ml. L-arabinose was added to the medium at a final concentration of 0.2% to induce the expression of genes cloned downstream of the araBAD promoter in vector pBAD33.

### DNA manipulations

Plasmid DNA was isolated using a Plasmid Mini Isolation Kit (A&A Biotechnology) according to the manufacturer’s instructions. DNA was introduced into bacterial cells by electroporation, using 1-mm gap cuvettes (BTX) and a MicroPulse electroporator (Bio-Rad), as described by Sambrook and Russell (2001). Details of plasmid constructions are presented in Supplementary Table S1. Routine DNA manipulations were carried out using standard procedures (Sambrook and Russell, 2001). All restriction, DNA-modifying enzymes, and DNA ligase were supplied by Thermo Fisher Scientific. Amplification of DNA fragments by PCR was performed using Pfu or Taq DNA polymerase (Thermo Fisher Scientific), appropriate primers, and template DNAs. Point mutations in the *repM* gene were generated using specific primers and a QuikChange Site-Directed Mutagenesis kit according to the protocol supplied by the manufacturer (Stratagene). All oligonucleotide primers used in this study are listed in Supplementary Table S2.

### Plasmid stability assay

Segregational stability of pIGMS31 plasmid derivatives was tested by replica plating following growth under non-selective conditions for approx. 80 generations, as described previously (Bartosik et al., 2016). The incompatibility characteristics of two plasmids (residing and introduced) were examined by testing the stability of the residing replicon (pMS-5) in the presence of ampicillin in the growth medium (antibiotic selection for introduced pUC18 vector derivatives containing putative determinants of incompatibility of pIGMS31).

### Overexpression and purification of 6His-tagged RepM protein

RepM protein was overexpressed and purified using a method described by Rozhon (2017) with some modifications. The *repM* gene pIGMS31 was cloned in expression vector pET28b+ resulting in pET-*repM* (for RepM(6His) overexpression) (Supplementary Table S1). *E. coli* BL21(DE3) strain harboring pET-*repM* was cultured overnight in LB medium supplemented with kanamycin at 37°C with shaking (180 rpm). For protein overexpression, 8 ml of the overnight cultures were added to 1000 ml of fresh LB + kanamycin medium, and incubation was continued at 28°C with shaking (180 rpm). When the culture had reached an OD_600_ of between 0.35 and 0.4, isopropyl *β*-D-1-thiogalactopyranoside (IPTG) was added to a final concentration of 0.4 mM to induce the expression of the 6His-tagged proteins. The cultures were then incubated further to an OD_600_ 1.0 under the same conditions. The cells were collected by centrifugation (15 min, 6,500 xg, 4°C) and resuspended in 15 ml of lysis buffer: 50 mM sodium phosphate pH 8.0, 300 mM NaCl, 10 mM imidazole, 0.1% triton X-100, and 300 μl of 100 mg/ml lysozyme, supplemented with 1 mM PMSF and Protease Inhibitor Cocktail (Sigma). After holding on ice for 15 min the cells were disrupted by sonication and the obtained lysate was centrifuged (15 min, 22,000 xg, 4°C) to pellet cell debris. All subsequent steps were performed at 4°C. The cleared lysate was incubated with 0.25 ml of Ni-NTA beads (Qiagen) for 30 min, with gentle shaking. The Ni-NTA resin was then given a series of washes: (i) twice with 8 ml of W1 buffer (50 mM sodium phosphate pH 8.0, 300 mM NaCl, 20 mM imidazole), (ii) once with 4 ml of W2 buffer (50 mM sodium phosphate pH 8.0, 2 M NaCl, 20 mM imidazole), and finally (iii) three times with 8 ml of W1 buffer. The 6His-tagged proteins were finally eluted with 0.4 ml of elution buffer (50 mM sodium phosphate pH 8.0, 300 mM NaCl, 150 mM imidazole). The concentration of the purified recombinant protein was estimated using the Bradford dye-binding method. Protein was analyzed by SDS-PAGE (Supplementary Figure S2) and MALDI-TOF mass spectrometry to confirm their identity. Protein aliquots were frozen in liquid nitrogen and stored at - 70°C.

### Electrophoretic mobility shift assay

#### Obtaining fluorescein (FAM)-labeled DNA fragments

The following elements of the REP module of pIGMS31 were cloned in vector pUC18: (i) CR (pUC-MS_13), (ii) IT1-2 (pUC-MS_17), IT3-4 (pUC-MS_18), (iii) IR1, IR2 (pUC-MS_15), (iv) IR2 (pUC-MS_14), and (v) *C*-terminal part of *repM* (pUC-MS_20) (Supplementary Table S1). Using these plasmid constructs as templates the cloned DNA fragments were amplified by PCR with “universal” M13 forward and reverse primers – M13pUCr and FAM-labeled M13pUCfFAM, oligos 59 and 60 (Supplementary Table S1), and subsequently purified using a Clean-Up kit (A&A Biotechnology). The same primer pair was used for the amplification of a 136-bp DNA fragment of pUC18, which served as a negative control.

#### DNA binding assay

Binding reactions of the total volume of 18 μl, containing 2 μl of 10x EMSA buffer [200 mM Tris-HCl (pH 8.0), 1000 mM KCl, 10 mM EDTA, 3 μg of poly-dIC competitor, 40 μg of BSA (bovine serum albumin)], were incubated for 5 minutes at room temperature with 1 μl of RepM(6His) (approx. 500 ng - 16 pmol) or H_2_O (control reaction). The approximate protein amounts were calculated based on the Bradford dye-binding method and SDS-PAGE analysis. Subsequently, 0.25 pmol of FAM-labeled DNA fragments were added to the final volume of 20 μl. After 30 min incubation at 25 °C, the reactions were gently mixed with 6 μl of 50% glycerol and loaded on a standard 6% polyacrylamide gel cast with TBE. Protein-DNA complexes were then separated by electrophoresis in 1 x TBE buffer at 10 V/cm at 10 °C and DNA fragments were visualized using Imager 600 (Amersham).

#### DNA relaxase activity assay

To determine if RepM has nuclease/relaxase activity, the procedure described by Rozhon and coworkers (2011) was adopted. 1 μg of supercoiled DNA (scDNA) of pIGMS31 derivative pMS-5 was incubated with increasing amounts of purified RepM(6His) protein in reaction buffer [25 mM Tris-HCl pH 7.6, 1 mM DTT, 0.1 mM EDTA, 5 mM Mn^2+^]. After incubation samples were treated with proteinase K and loaded on 1% agarose gel, and electrophoresed. DNA was visualized by Gel-Red staining.

### 5′ RACE for determination of the *repM* transcriptional start site

#### RNA isolation

An overnight bacterial culture was diluted 1:50 in fresh medium and cultivated for a further 3 hours. The cells were harvested by centrifugation and total RNA was isolated from the pellet using a Total RNA Mini Plus kit (A&A Biotechnology) and DNase was treated using a Clean-Up RNA Concentrator kit (A&A Biotechnology). The quality and concentration of the isolated RNA were evaluated using a Picodrop Microliter UV/Vis Spectrophotometer and standard formaldehyde agarose gel electrophoresis. RNA was stored at -20°C.

#### 5’ RACE (rapid amplification of cDNA ends)

To determine the *repM* transcription start site a 2nd Generation 5’/3’ RACE Kit (Roche) was used according to the manufacturer’s instructions with some modifications. Since the pIGMS31 sequence contains a high level of A+T pairs, Poly[C] tailing of cDNA was applied instead of Poly[A] tailing. Consequently, the Oligo dT-anchor primer (included in the kit) was replaced by an Oligo dG-anchor primer, oligo 64 (Supplementary Table S2), during the primary PCR. For cDNA synthesis, 1 μg of total RNA was used with the gene-specific primer SP1RACEMS, oligo 65 (Supplementary Table S2). Two separate PCR amplifications were performed using dG-tailed cDNA as the template, the first with an Oligo dG-anchor primer (included in the kit) and gene-specific primer SP2RACEMS, oligo 65 (Supplementary Table S2). The products of these primary reactions were then used as the template in a secondary PCR, with PCR anchor primer (included in the kit) and gene-specific primer SP3RACEMS, oligo 66 (Supplementary Table S2). The amplified DNA fragment was visualized by agarose gel electrophoresis, then purified using a Gel-Out kit (A&A Biotechnology) and sequenced.

#### 5′ RACE determination of RepM nicking site

The 5’ RACE method was used to determine the exact position of DNA cleavage performed by RepM within the pIGMS31 replication origin. A template for the C-tailing reaction was prepared as described above in the DNA relaxase activity assay section with some modifications. After incubation of plasmid scDNA with RepM(6His) protein 5’ RACE template was purified by phenol-chloroform extraction and ethanol precipitation. Dissolved DNA was incubated with terminal transferase (TdT) for C-tailing of the 3’ end of the nicked DNA strand. Next, two PCR reactions were performed with PCR anchor primer (included in the 2nd Generation 5’/3’ RACE Kit, Roche) and GB3 (oligo 61) or GB9 (oligo 62) (Supplementary Table S2). In this way, we checked the possibility that nick can be introduced on the + or - DNA strand. Only in the case of reaction with GB3 primer PCR product (single band) was obtained. After electrophoresis DNA was purified and AT cloned into a pDRIVE vector. Plasmid DNA of four separate clones harboring recombinant pDRIVE with cloned 5’ RACE product was sequenced giving the same result of nick mapping.

#### *In vivo* RepM protein interactions assay

RepM protein interactions were analyzed using a bacterial two-hybrid system constructed by Di Lallo and co-workers (2001). This system uses the *E. coli* R721 test strain carrying chromosomally encoded *lacZ* reporter gene and two expression vectors (both enabling recombinant proteins production from IPTG inducted *lacZ* promoter). The *lacZ* promoter present in the *E. coli* R721 chromosome contains a hybrid operator with binding sites for (i) phage P434 repressor (encoded in the test vector pcl434) and (ii) phage P22 repressor (encoded in the test vector pcl22). If the analyzed proteins, overproduced in translational fusions with the repressor subunits, interact directly with each other, an active hybrid repressor is formed and R721 *lacZ* promoter activity is inhibited. The measure of dimer formation in this system is therefore the decrease in *β*-galactosidase activity in comparison with the control strain R721. Descriptions of the constructed recombinant pcl22 and pcl434 vectors are listed in (Supplementary Table S1).

#### *β*-galactosidase assay

*β*-galactosidase activity assays were performed according to a method described by Thibodeau and co-workers (2004), with slight modifications. *E. coli* strains were cultivated overnight in a liquid LB medium supplemented with suitable antibiotics. These cultures were diluted 1:50 in fresh medium and cultivated for a further 2 hours. For the Rep protein interaction assay (bacterial two-hybrid system) the LB medium was supplemented with IPTG to a final concentration of 0.1 mM according to Di Lallo and co-workers (2001). The OD_595_ of the bacterial cultures was then measured and eight replicates of 80 μl of each culture were transferred to a 96-well microtiter plate. To cause cell lysis, 20 μl of PopCulture lysis buffer (Merck Millipore) was added to each well, and the mixtures were incubated for 15 min at room temperature. Twenty microliters of the lysates were then added to wells of another 96-well plate containing 130 μl of Z-buffer. To initiate the enzymatic reaction, 30 μl of the *β*-galactosidase substrate ONPG (4 mg/ml) (Sigma) was added to each well. The plate was then placed in a TECAN plate reader (Tecan Group Ltd) and incubated at 28°C as the OD_415_ was measured at 30 s intervals.

#### Replication intermediates analysis by two-dimensional gel electrophoresis

Isolation of DNA replication intermediates (RIs) and two-dimensional neutral/neutral DNA agarose gel electrophoresis (2D gel) was performed according to Schvartzman and co-workers (2012). Non-radioactive hybridization was performed according to the mentioned above paper (Schvartzman et al., 2012) with modifications - membranes were pre-hybridized in a volume of 0.2ml of pre-hybridization solution (5xSSC, 0.1% N-Lauroylsarcosine sodium salt, 1% SDS, 2% Blocking reagent (Roche)) for each 1cm^2^ of the membrane. Prehybridization lasted at 55°C – 65°C for 2–4 h. Hybridization probe was obtained using DIG High Prime DNA Labeling Kit (Roche) and linearized pMS-7 DNA as a template. Then labeled DNA was added and the subsequent stages of the procedure were continued according to Schvartzman and co-workers (2012).

#### Bioinformatic analyses

DNA or protein sequences were aligned using BLAST (https://blast.ncbi.nlm.nih.gov/Blast.cgi) (Altschul et al., 1997). Protein secondary structures were predicted with YASPIN Secondary Structure Prediction (https://ibi.vu.nl/programs/yaspin) (Lin et al., 2005). Molecular masses and isoelectric points of proteins were predicted using Compute pI/Mw from the Expasy server (https://web.expasy.org) (Artimo et al., 2012).

## Results

### Genetic organization of pIGMS31 plasmid

pIGMS31 is a small 2520 bp plasmid composed of two genetic modules, responsible for the initiation of replication (REP) and mobilization for conjugal transfer (MOB) (Figure 1A, Supplementary Figure S3) (Smorawińska et al., 2012). It encodes only two ORFs, *repM* (in REP module) for RepM protein (201 aa) and *mobM* (in the MOB module) for MobM protein (390 aa). MobM protein has a typical amino acid signature of the MOB_V_ family mobilization proteins [Motif I (H-x2-R) and Motif III (H-x-DE-x2-PH-x-H)] (Francia et al., 2004). RepM shares sequence similarity to atypical replication initiators of the pHW126-like plasmids family. It was proposed that they are rolling circle (RC) replicons, but their Rep proteins, including RepM, do not contain sequence motifs characteristic for RC replication initiators of the HUH or *Rep_trans* families (Wawrzyniak et al., 2017). Using the BLAST server, a helix-turn-helix (HTH) motif was identified in the central part of the pIGMS31 RepM protein. The predicted secondary structure of aa sequences surrounding this putative HTH motif resembles wHTH (winged helix-turn-helix) DNA binding/dimerization motifs of MarR (multiple antibiotic resistances regulator)-like transcriptional regulators (Lin et al., 2005; Perera and Grove, 2010) (Supplementary Figure 4). Also, putative components characteristic of pHW126-like REP modules were identified within the REP module of pIGMS31. They are: (i) conserved region (CR), (ii) four iteron-like 20 bp direct repeats (IT1-4), (iii) a palindromic sequence similar to single-strand initiation sites for priming DNA replication (*ssi*) found in diverse replicons (Nomura et al., 1991). In addition, *repM* gene is directly preceded by two copies of short 9 bp inverted repeats IR1 and IR2.

**Figure 1.**
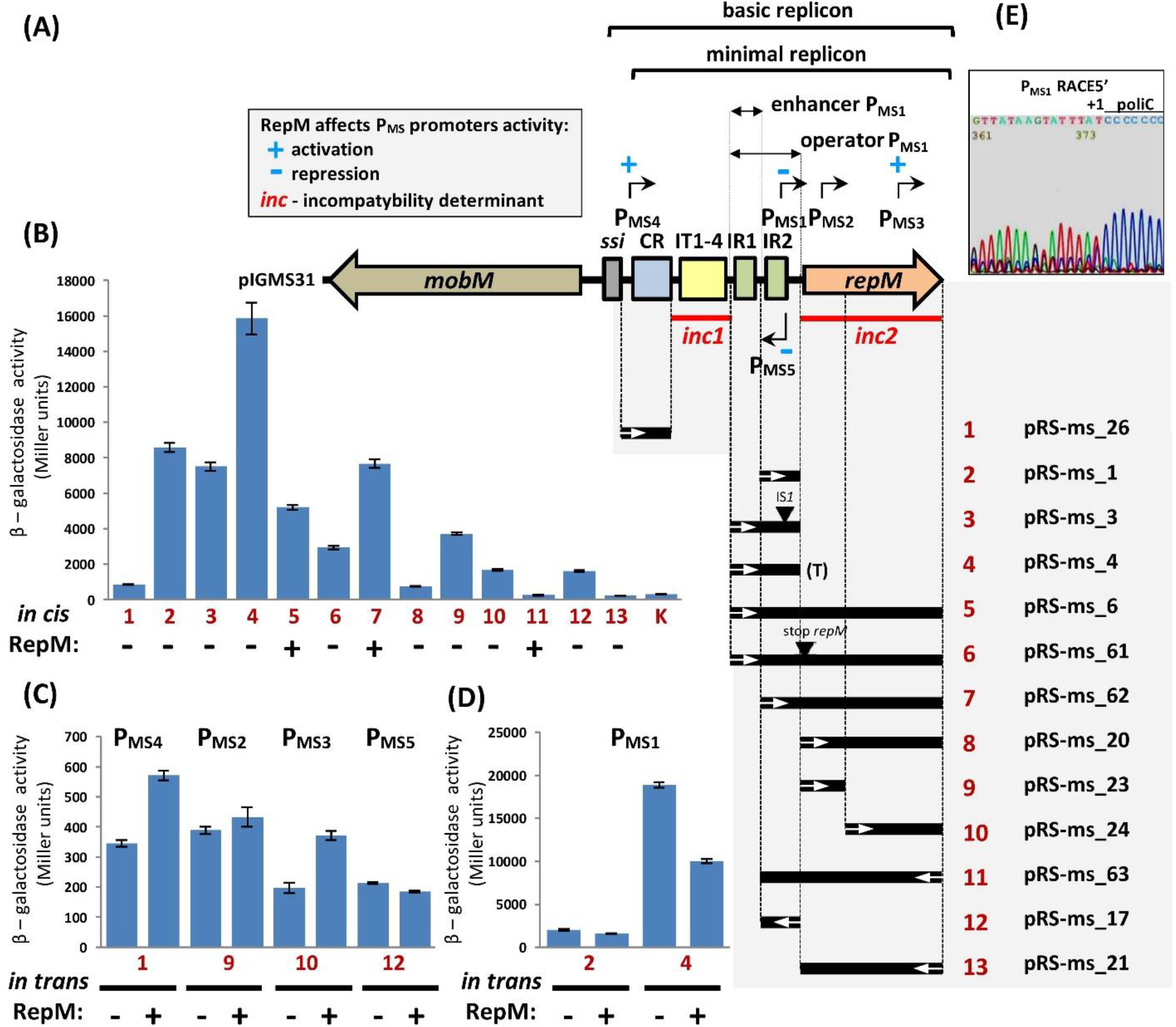
Functional analysis of the replication system of plasmid pIGMS31. **(A)** Schematic representation of plasmid genome. Elements of basic and minimal replicon are indicated: single-strand initiation site (*ssi*), conserved region (CR), iteron-like sequences (IT), and inverted repeats (IR). Promoters identified within the REP module are designed as black arrows. Operator and enhancer elements of the P_MS1_ promoter are indicated. For more detail see Supplementary Figure 4-6. Analysis of P_MS_ promoters activity and their regulation: black lines represent DNA fragments of pIGMS31 and their mutated versions, and white arrows set the orientation of DNA fragments cloned in the pRS551 test vector. Black triangles mark the IS*1* integration site and mutation of *repM* ATG start codon to stop codon. (T) – T*pro*/T*lyz* transcriptional terminator of P1 phage. **(B)** *β*-galactosidase activity of protein extracts from strains carrying pRS551-derived constructs imaged in panel (A), reflecting the strength of P_MS_ promoters, K – “empty” test vector. **(C, D)** P_MS_ promoter activity in the presence or absence of RepM protein derived *in trans* from pBAD vector **(E)** Sequencing result for the *repM* 5’ RACE product. The chromatogram represents the *repM* coding strand. The *in vitro* synthesized poliC tail is indicated and the predicted *repM* transcription start site is marked by +1.

### Identification of pIGMS31 basic and minimal replicon

To delineate a minimal DNA region of pIGMS31 capable of autonomous replication, several deletion derivatives of this plasmid were constructed and their replication abilities were tested in *E. coli* cells (Supplementary Figure 5). This analysis revealed that all but one (*ssi*) of the aforementioned REP components are necessary for replication. The presence of *ssi* was not obligatory, although its absence caused an increase in copy number of the constructed plasmid and its rapid multimerization, which greatly reduced the stability of this replicon in bacterial cells (Supplementary Figure S5).

The *origin* of pIGMS31 replication was defined using a two-plasmids system described previously (Wawrzyniak et al., 2019). Selected parts of the REP region were amplified by PCR and ligated to a DNA cassette containing a kanamycin resistance gene (KM) and *oriγ* of plasmid R6K (the presence of *oriγ* enabled replication of the constructed plasmids in *E. coli* DH5αλpir, providing the π protein of plasmid R6K that initiates replication at *oriγ*). The constructed plasmids were then introduced into *E. coli* DH5α (lacks the *pir* gene) carrying plasmid pUC-*repM*_1, as a source of the pIGMS31 RepM protein. The introduced plasmids were able to replicate only when they contained a functional pIGMS31 *origin*, activated by the *trans-*acting RepM protein. This analysis revealed that the minimal *origin* is included in 258 bp pIGMS31 DNA fragment containing CR, IT1-4, and IR1 sequences cloned in construct pMS-6γ (Supplementary Figure S5). In the next step, the incompatibility properties of predicted REP elements were tested. For this purpose, different REP fragments were cloned into pUC18 (compatible with pIGMS31). Obtained constructs were introduced into an *E. coli* strain containing plasmid pMS-5 (Km resistant derivative of pIGMS31). Removal of the residing parental replicon was indicated in the case of pUC18 vectors harboring IT1-2, IT3-4 sequences, and complete *repM* ORF. In this way two incompatibility regions were distinguished *inc1* (IT1-4) and *inc2* (*repM* ORF) respectively. Details of these analyses of the pIGMS31 REP module are presented in Supplementary Figure 6 and summarized in Figure 1A.

### Mapping of *repM* gene promoter

To define mechanisms responsible for the regulation of the *repM* gene expression, precise location of the *repM* transcription start site (+1) was mapped by 5’ RACE. It was located at position 16 of the pIGMS31 sequence (Figure 1E) within the right arm of the IR2 sequence. Notably, the nearby hexameric sequences [5’-TTGACT(15N)TATAAC-3’] are almost identical to the *E. coli* promoter consensus sequence [5’-TTGACA(15-19N)TATAAT-3’] (Shimada et al., 2014) (Fig. 1A).

A DNA fragment between the IT1-4 repeats and the start codon of *repM* (containing the predicted P_MS1_ promoter) was then cloned into test vector pRS551, to generate a putative transcriptional fusion with a promoter-less *lacZ* reporter gene (plasmid pRS-ms_3; Figure 1A). The level of *β*-galactosidase activity was then measured in a lysate of cells carrying this construct. Unexpectedly, just an average level of promoter activity was detected in *E. coli* DH5αΔ*lac* harboring pRS-ms_3 (Figure. 1B). Sequencing of plasmid DNA isolated directly from the bacterial culture used for the enzymatic assay showed that the P_MS1_ promoter had been partly inactivated by transposition of insertion sequence IS*1* (Figure 1B). Analogous transposition events were observed in two other independent experimental approaches (data not shown). It is assumed that the genetic instability of pRS-ms_3 might result from the extremely high activity of the native P_MS1_ promoter, deprived of its regulatory elements (as was demonstrated in the case of *repR* gene promoter from pIGMS31 related plasmid pIGRK (Wawrzyniak et al., 2019)).

To verify this hypothesis, a strong transcription termination signal, derived from P1 prophage (T*pro*/T*lyz* terminator (Bartosik et al., 2012)), was inserted into pRS-ms_3, between the predicted P_MS1_ and the reporter gene. Diminished transcription from P_MS1_ stabilized the genetic structure of pRS-ms_4 (no IS*1* insertion mutants were selected). Moreover, despite the presence of a transcriptional terminator, more than two-fold increase in promoter activity was observed compared to pRS-ms_3 (Figure 1B). Interestingly, the decrease in P_MS1_ promoter activity (determined by IS*1* insertion) did not happen in the case of pRS-ms_1 (with P_MS1_ promoter deprived of IR1 sequence). It was also shown that the level of promoter activity cloned in pRS-ms_1 was significantly lower than in pRS-ms_4 suggesting a transcription enhancer function of sequence containing IR1.

### Identification of additional promotes within pIGMS31 REP module

To examine the autoregulatory role of the RepM protein towards *repM* gene expression, DNA fragments from pIGMS31 containing different variants of P_MS1_ were cloned in pRS551 test vector, along with the wild-type *repM* gene or the mutant *repM*stop (ATG → TAG substitution) (Figure 1A). *lacZ* reporter gene expression analysis using pRS-ms_6 vs pRS-ms_4 revealed that P_MS1_ promoter activity level decreases in the presence of *repM* (Figure. 1B) This observation lead us to hypothesize that RepM acts as a *repM* repressor. However, this regulatory effect was not confirmed mutating of *repM* start codon (ATG → TAG substitution) in pRS-ms_61. The lack of RepM did not result in increased activity but even in a decrease of the reported promoter activity (Figure 1B). More detailed *in silico* analyses, using BPROM, revealed the presence of two putative promoters within *repM* ORF with the same orientation as P_MS1_. Promoter activity was detected in the case of complete *repM* ORF (pRS-ms_20) or its fragments *repM*’ (pRS-ms_23) and ‘*repM* (pRS-ms_24). In this way, two additional promoters were identified: P_MS3_, in the *N*-terminal part of *repM*, and P_MS4_, in the *C*-terminal part of *repM* (Figure 1B). In our last paper, we proved that *repR* gene in pIGRK plasmid encodes two in-frame polypeptides: the replication initiator RepR and the negative regulator of replication RepR’(Wawrzyniak et al., 2019). Both RepR and RepR’ are also negative regulators of *repR* transcription (Wawrzyniak et al., 2019). However, instead of the GTG internal start codon of RepR’ there is a GTT codon in *repM* gene. We were not able to detect additional protein product(s) coded by *repM* by SDS-PAGE-Western blot, nor mutations of putative internal start codons (data not shown). on the other hand, *in silico* search for promoters within the pIGMS31 REP module found two more promoters. One, named P_MS4_, is located within the minimal replication *origin* near the CR element. The second, named P_MS5_, is located within the P_MS1_ coding sequence (but it is positioned in opposite direction) (Figure. 1A). Transcriptional activity promoted by DNA fragments containing P_MS4_ and P_MS5_ was confirmed by cloning them into vector pRS551 followed by *β*-galactosidase activity measurement (Figure 1B).

### Examination of RepM transcriptional regulator properties

We decided to investigate the role of RepM in the regulation of each P_MS_ promoter activity using *in vivo* two-plasmids assay. Test strains contained pRS551 plasmid, with cloned pIGMS31 REP promoter sequences, and plasmid pBAD33-*repM* (source of RepM protein) or empty pBAD33 (used as a control). Promoter activity was measured in cell lysates of bacteria cultivated in LB medium supplemented with the appropriate antibiotics and with 0.2% arabinose for the induction of araBAD promoter. In this experiment, it was demonstrated that the activity of pIGMS31 REP promoters, except P_MS2_, was affected by *trans*-expression of RepM protein (Figure 1CD). P_MS1_ activity was down-regulated by RepM especially, when P_MS1_ was cloned with the IR1 sequence. Also, P_MS5_ activity was lowered by the presence of RepM. However, this effect was more evident when *repM* gene was present in cis (promoter activity of DNA fragment cloned in pRS-ms_63 vs pRS-ms_17) (Figure 1BC). In contrast, P_MS3_ and P_MS4_ promoters were activated in the presence of RepM protein (Figure. 1C).

### RepM DNA-binding properties

Direct interactions of RepM with DNA fragments from pIGMS31 REP were investigated by electrophoretic mobility shift assays (EMSA). Recombinant RepM(6His) was purified and mixed with FAM-labeled DNA fragments containing elements indispensable in cis for plasmid replication (Figure 2A): CR (also containing P_MS4_), IT1-4, the potential operator sites of the P_MS1/5_ promoters region (IR1, IR2), and the *C*-terminal part of *repM* gene (containing P_MS3_), as well control fragment of thefrom pUC18 plasmid (K). The highest DNA binding capacity of RepM(6His) was detected with fragments 4 and 5, containing IR1 and IR2 or only IR2 sequences, respectively (Figure 2B). Most of the molecules of DNA fragments 4 and 5 were retarded as nucleoprotein complexes with RepM(6His). Four shifts were formed by fragment 4 and two by fragment 5. In the case of the other pIGMS31 REP DNA fragments used, significantly less abundant complexes were detected as double shifts. This applies to both minimal replication *origin* elements, fragments 1 (CR with P_MS4_), 2 and 3 (IT1-2 and IT3-4 respectively), and to fragment 6 (containing P_MS3_). The presence of a single shift was also formed in the case of control fragment K (Figure 2B). It can be interpreted as a demonstration of residual nonspecific RepM DNA binding properties.

**Figure 2.**
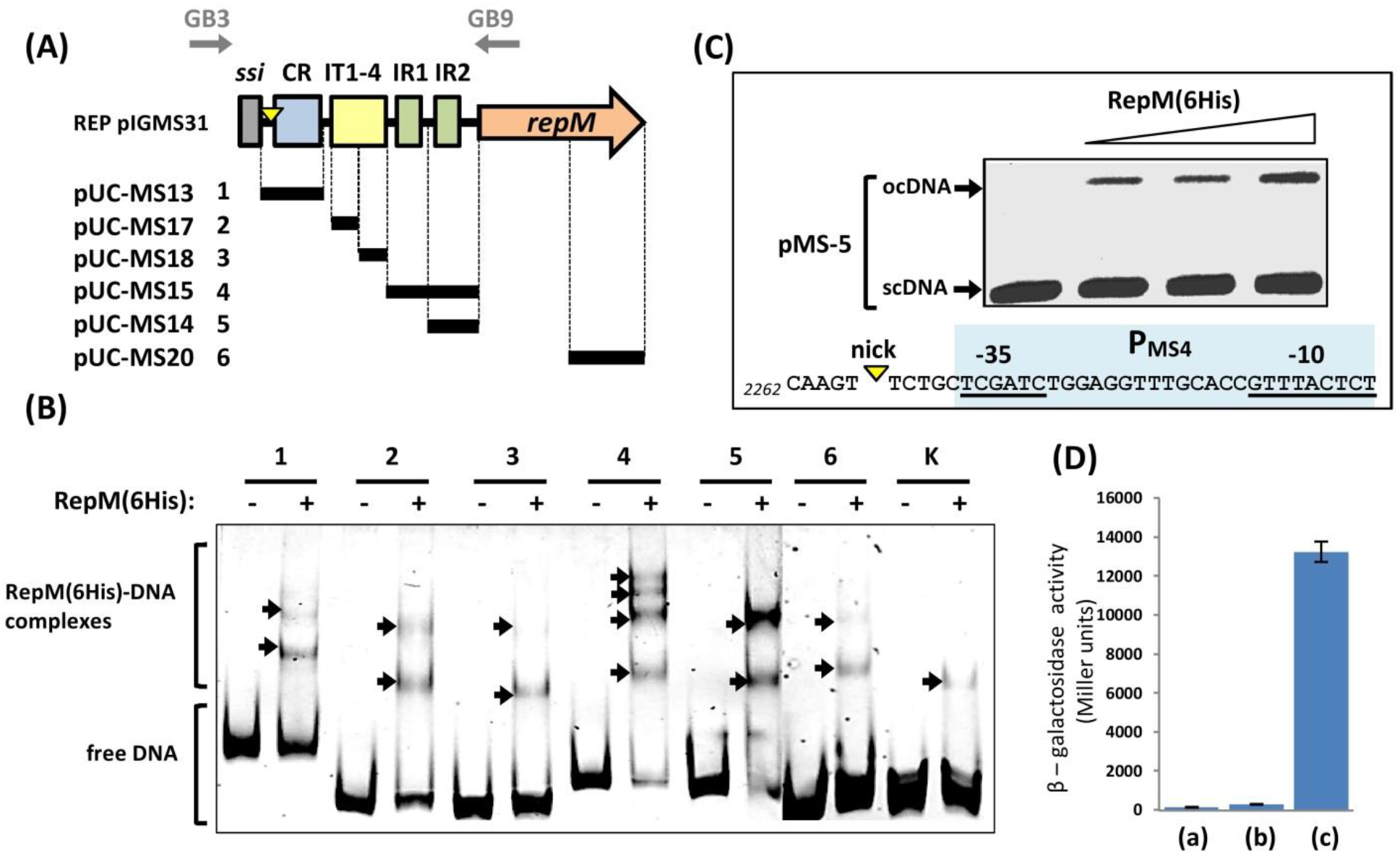
Functional characterization of the RepM protein. **(A)** DNA fragments from pIGMS31 REP cloned in pUC18 plasmid used in EMSA experiment. GB3 and GB9 correspond to the primers used for 5’ RACE detection of the nick introduced by RepM within pIGMS31 replication *origin*. **(B)** Electrophoretic mobility shift assay (EMSA). Binding of RepM(6His) to FAM-labeled DNA fragments of pIGMS31 presented in panel (A). Nucleoprotein complexes are marked with black arrows. In lane K – the control DNA fragment from plasmid pUC18 was used. **(C)** Determination of RepM DNA relaxase activity. Supercoiled (sc) DNA of pMS-5 was incubated with increasing amounts of recombinant RepM(6His) protein. In the presence of recombinant RepM, plasmid scDNA was converted to relaxed open circle (oc) DNA. **(D)** Results of RepM dimerization properties measurement by the bacterial two-hybrid system. Decrease of *β*-galactosidase activity is a measure of dimer formation by: (a) RepM, (b) SXT proteins – as positive control (Dziewit et al., 2007), and (c) negative control.

### RepM forms dimers *in vivo*

Most bacterial plasmids Rep proteins exist as a dimer. Interestingly RepM protein of pIGMS31 differs from pHW126 and pIGRK and Rep proteins in absence of a putative *C*-terminal dimerization coiled-coil motif (Wawrzyniak et al., 2019). To test the ability of RepM to form homodimers, a bacterial two-hybrid system was used. *repM* gene was cloned into the test vectors pcl434 and plc22 (Figure 2D) to produce plasmids able to express recombinant RepM (pcl434M and plc22M) proteins. Analysis using these constructs showed that RepM molecules interact with each other to form homodimers (Figure 2D).

### RepM has DNA relaxase activity

A typical feature of RC plasmid replication initiators is DNA relaxase activity. RC Rep protein introduces a site and strand-specific break (nick). After cleavage reaction, Rep remains attached to the 5’ end of the break while the free 3’-OH end is used as a primer for lagging strand synthesis (Wawrzyniak et al., 2017). This type of nuclease activity was demonstrated for the Rep protein of pHW126 plasmid (Rozhon et al., 2011). Using the same experimental procedure (as for Rep from pHW126), RepM relaxase activity was examined. Supercoiled DNA of pMS-5 preps was incubated with increasing amounts of RepM(6His) protein. After incubation samples were electrophoresed and visualized (Figure 2C). In the sample incubated without RepM, only a single band corresponding to the supercoiled pMS-5 DNA was visible. In samples containing recombinant RepM, an additional signal corresponding to relaxed open-circle DNA was detected.

Next, we determine the DNA cleavage site iusing the 5’ RACE’ technique. The usefulness of this method was validated by the determination of a specific unique DNA nick introduced by NbBpu10I within the pBGS18 plasmid (data not shown). The 3’-OH end of nicked DNA was poli-C-tailed by terminal transferase and used as a template in PCR reactions with one of two plasmid-specific primers GB2 and GB9 spanning the predicted minimal replication origin (Figure 2A). In this way, we were able to detect nicks on both DNA strands. Only in the case of amplification with GB3 primer a PCR product of about 500 bp was obtained. DNA sequencing of the PCR product (AT cloned in pDrive vector) revealed the location of DNA processing between positions 2268 and 2269 on the + strand, just upstream the *in silico* predicted P_MS4_ promoter -35 hexamer sequence (Figure 2A). In a control experiment using a substrate supercoiled pUC18 plasmid (lacking the pIGMS31 REP module), similar relaxase activity was demonstrated. In this case, multiple nicking sites were determined within pUC18 DNA treated with RepM(6His), suggesting that RepM may act in a non-specific way in the absence of a cognate target sequence (data not shown).

### Analysis of DNA RIs by two-dimensional gel electrophoresis (2D gels)

Determination of DNA relaxase activity of Rep protein is believed as a “gold standard” in RC bacterial plasmids identification (del Solar et al., 1998). Using this criterion it was proposed that pHW126, and other related plasmids, represent the new RC replicons family. On the other hand, our findings demonstrate that pIGMS31, apart from RepM nuclease activity, exhibits many features not reported for known RC plasmids. In some cases they are features customarily associated with *theta* plasmids. IT1-4 iteron-like sequences are present in the replication origin acting as RepM binding sites and incompatibility determinants; RepM protein contains a wHTH motif ;and RepM shows autoregulatory properties interacting directly with its gene promoter. For this reason, we decided to study the replication mode of pIGMS31 by analyzing the replication intermediates (RIs) that are generated *in vivo*. We used two-dimensional (2D) agarose electrophoresis, a technique widely used for this purpose, reviewed in Schvartzman et al. (2010)

Briefly, plasmid DNA from cells growing in exponential phase is purified under neutral pH conditions and digested with restriction enzymes to generate the linear DNA fragments to be studied. Next, the DNA is separated by 2D electrophoresis in agarose gals and a Southern blot is performed hybridizing with probes specific for the fragments under study. Electrophoresis conditions during the first dimension allow molecules to be separated based mainly on their size (mass). On the other hand, the conditions during the second electrophoresis cause the shape of the molecules to significantly affect mobility, such that branched replication intermediates or those with an internal bubble suffer a delay compared to their behavior in the first electrophoresis. In this way, the hybridization pattern observed after hybridization allows knowing the replication mode of the fragment under study. The basic patterns that can be observed in the case of *theta* type replication are shown in Figure 3A.

**Figure 3.**
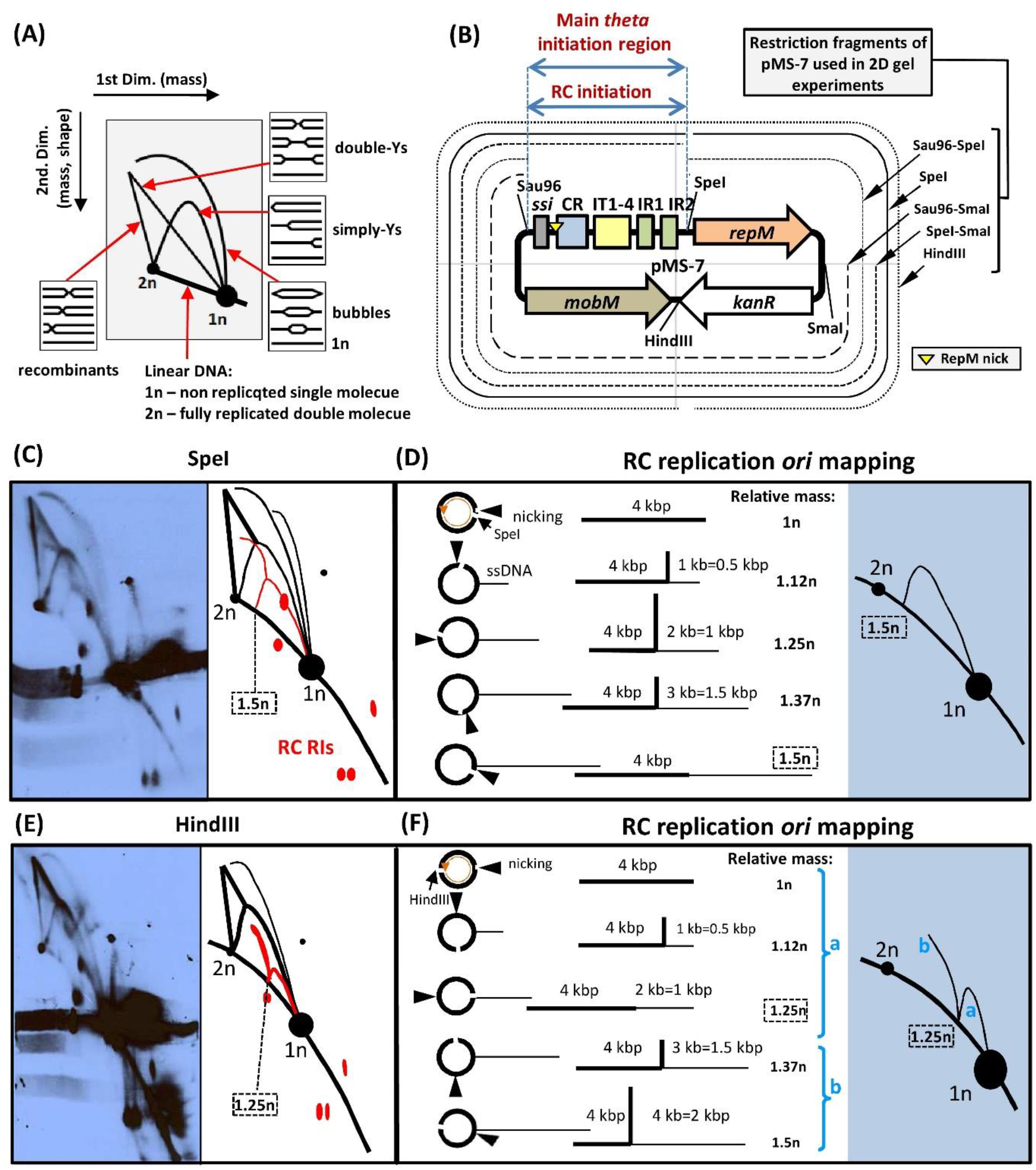
2D gel analysis of pMS-7 RIs with explanatory drawings. **(A)** Scheme representing the shape of the *theta* replication intermediates and the basic 2D patterns generated. **(B)** Diagram representing the genetic map of pMS-7 plasmid and the restriction fragments analyzed in 2D gels (analysis of HindIII and SpeI fragments are shown below). Plsmid region where it is suggested initiation of RC and *theta* replication takes place is indicated. **(C and E)** 2D gel immunograms and summary drawings showing major replication intermediates inferred from the gels upon linearization of plasmid pMS-7 with SpeI (C) or HindIII (E). Signals generated by RIs arising from RC replication are depicted in red. **(D and F)** Cartoon of the expected RC replication intermediates before and after plasmid linearization with SpeI (D) or HindIII (F). Thin lines correspond to ssDNA. To the right, the expected 2D patterns are shown.

Our first 2D electrophoresis of pIGMS31 derivatives generated complex hybridization patterns that were difficult to interpret, probably due to high multimer formation and weak signal intensity. Therefore, we used pMS-7, a spontaneous high copy number mutant of pMS-5 that carries a single residue substitution in the *repM* sequence (471G>C), resulting in the RepM 140A>P mutation (Figure 3).

We have analyzed by 2D electrophoresis a series of overlapping fragments of pMS-7 after their digestion with the restriction enzymes indicated in Figure 3B. Shown here are the results of the analysis of replication intermediates observed after linearization of the plasmid with SpeI or HindIII. In plasmid pMS-7 linearized with SpeI, the *origin* of replication containing CR and IT1-4 elements is located centrally within the linear molecule. However, by using HindIII to linearize the plasmid, the *origin* lies close to one end of the molecule. The expected rolling circle replication intermediates when pMS-7 is linearized with SpeI are diagrammed in Figure 3D. The non-replicating linear plasmid is about 4 kb (relative mass=1n) and from which a strand of ssDNA is generated by strand displacement. The size of this ssDNA branch increases progressively as the synthesis of the (+) strand DNA advances and its completion. This latter intermediate is linear and its relative mass is 1.5n, as half of the molecule is single-stranded. As shown in the diagram of Figure 3D (right panel), it is expected these RIs generate in the 2D gel an arc that starts at 1n, inflects, and falls to a position corresponding to a linear molecule of relative mass 1.5n. This pattern expected in the case of RC replication was indeed visible in the 2D gel (Figure 3C). Similarly, the schem of Figure 3F shows the expected RIs after linearization of pMS-7 with HindIII. The expected pattern generated by these RIs (Figure 3F, right panel) was also visible in the 2D electrophoresis (Figure 3E).

These results indicate that PMS-7 is capable of rolling circle replication and that initiation occurs in the region where CR and IT1-4 elements are located, between SpeI and Sau3A restriction sites (Figure 3B). This is consistent with previous results from *in vitro* RepM nicking site determination (Figure 2).

However, in addition to the signals described above, other patterns were also observed that are not expected in the case pMS-7 replicated exclusively by rolling circle (Figure 3C and E). These signals include a full arc corresponding to intermediaries containing a *theta*-mode replication initiation bubble, and a full arc corresponding to Y-shaped RIs, generated by a theta-type replication fork that runs through it. The presence of these patterns in the 2D gels indicates that pMS-7 also replicates in a *theta*-like pattern. This *theta* replication probably starts more often, but not only, in the same region where rolling circle replication initiates.

## Discussion

In this work, we performed a compressive study on the REP module of pIGMS31 plasmid from *K. pneumoniae*. Individual elements of the pIGMS31 replication system were identified and characterized in detail. In addition, initial studies on pIGMS31 replication mode were performed. Our work makes a significant contribution to the knowledge about the unique pHW126-like plasmids family. What is more, some general conclusions concerning bacterial plasmid replication have been drawn.

Until now, only two representatives of pHW126-like replicons were characterized. They are pHW126, characterized by Rozhon and co-workers (2010; 2011; 2012; 2017), and pIGRK described in our last publication (Wawrzyniak et al., 2019). Comparative analysis of REP modules from pIGMS31 as well as pHW126, and pIGRK revealed the presence of common core elements (Figure 4). They are the conserved region CR, and four iteron-like sequences IT1-4 present in their minimal replication *origin* as well as *in-trans* acting *rep* gene encoding replication initiation factor Rep protein composed of variable *N*-terminal sequence and most conserved wHTH motif (Supplementary Figure S4). Rep proteins are also acting as auto regulators and they inhibit cognate gene expression (Rozhon, 2017; Wawrzyniak et a., 2019), a typical feature of pHW126-like *rep* gene promoters. They show a very high transcriptional activity (especially in the case of *rep* promoters from pIGMS31 and pIGRK) and complex structure containing operatory (DR and/or IR elements) and enhancer sequences (Rozhon, 2017; Wawrzyniak et a., 2019 and this study). Whole three plasmids contain also an additional (not essential for replication) element, *ssi*-like palindromic sequence. This type of sequence is responsible for primosome complex formation and initiation of lagging strand replication in some *theta* and RC plasmids (Nomura et al., 1991; Ruiz-Masó et al., 2015). What is unique and typical for these pHW126-like replicons, is that the lack of *ssi* results in plasmid multimer formation (Rozhon et al., 2012; Wawrzyniak et a., 2019 and this study). This feature of *ssi* was not reported in previous studies on plasmid or phage replication systems containing *ssi* signals (Rozhon et al., 2012). Our investigations revealed additional common components present (at least) in REP modules of both pIGMS31 and pIGRK, as well as specific elements present only in pIGMS31 or pIGRK (Figure 4). Furthermore, we demonstrated that even core elements of REP from pIGMS31, pIGRK and pHW126 varied structurally and functionally. In this way, we proved that REP modules of pHW126-like plasmids are more complex and more diverse than would be expected based on comparative DNA sequence analyses.

**Figure 4.**
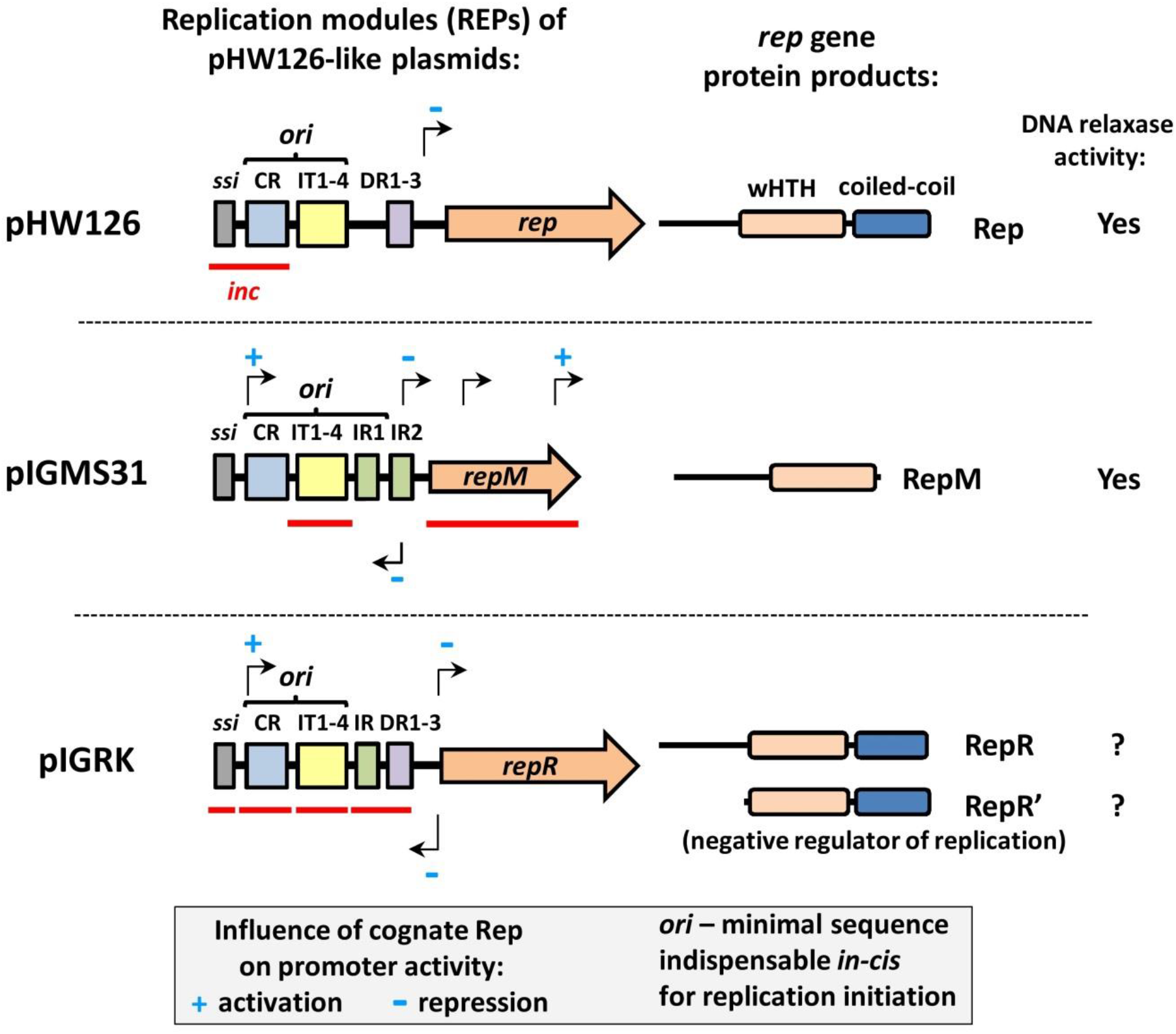
Schematic representation of REP modules encoded by pHW126-like replicons

In pIGMS31 four additional promoters were identified within the REP module (Figure 1). Equivalents of two of these promoters were identified also in the pIGRK plasmid (for more detail see Supplementary Figure S7). P_MS4_ (pIGMS31) and P_RK3_ (pIGRK) promoters are located within *in cis*-acting minimal replication *origins*. Both are positively regulated in the presence of cognate replication initiators RepM and RepR, respectively. It suggests that P_MS4_ and P_RK3_ activity may support replication initiation in some way. We were not able to identify potential genes (and their protein products) downstream from P_MS4_ and P_RK3_. We speculate that P_MS4_ and P_RK3_ may be involved in transcriptional activation of the replication *origin*, which was proposed for λ phage (Węgrzyn and Węgrzyn, 1995) and RC plasmid mp1 (Backert, 2002) or in RNA primer synthesis e.g. in *theta* ColE1 plasmids (Lilly et al., 2015). Two other promoters, P_MS5_ (pIGMS31) and P_RK2_ (pIGRK) overlap with promoters of *rep* genes P_MS1_ (pIGMS31) and P_RK1_ (pIGRK) but they are divergently oriented (towards replication *origins*). In contrast to P_MS4_ and P_RK3_ promoters, P_MS5_ and P_RK2_ are negatively regulated by RepM and RepR respectively. Based on our results it is hard to determine the exact role played by P_MS5_ and P_RK2_ in replication initiation but it seems that they may act as antagonists of P_MS4_ and P_RK3_ respectively. In pIGMS31 two additional REP promoters were exclusively identified. Both P_MS2_ and P_MS3_ are positioned within *repM* ORF in the same orientation as *repM* promoter P_MS1_. Only P_MS3_ is regulated (activated) by RepM protein. The detection of P_MS2_ and P_MS3_ was unexpected since we failed to identify additional protein products of *repM* as we demonstrated in the pIGRK plasmid (Wawrzyniak et al., 2019). In *repR* gene, an additional internal translation start site was discovered for the RepR’ negative replication regulator. Despite this, no internal *repR* promoter activity was identified (Wawrzyniak et al., 2019; Supplementary Figure S7), even in the presence of RepR and/or RepR’ derived *in trans* (data not shown). However, we cannot exclude that *repM* encodes additional protein/RNA products and that its expression is regulated by internal *repM* promoter(s). It is more likely that complete *repM* gene ORF act as an incompatibility (*inc*) determinant (named *inc2*) - plasmid sequences directly involved in the regulation of REP module function (Figure 4). A second pIGMS31 *inc* region (named *inc1*) contains IT1-4 sequences. It is quite surprising since *inc* properties were reported only for iterons from *theta* plasmids (Chattoraj, 2000). In pIGRK, *inc* properties were indicated for IT1-4, but not for *repR* ORF. However, it must be noted that the negative regulator encoded by *repR*, RepR’ is expressed only from a promoter located outside *repR* ORF (RepR’ is not synthesized if only *repR* ORF is cloned) (Wawrzyniak et al., 2019). In contrast to pIGMS31, three additional *inc* regions are located within pIGRK REP, they are *ssi*, CR sequence, and the *repR* promoter operatory/enhancer sequences IR and DR1-3 (for more detail see Supplementary Figure S7). Finally, in the case of pHW126, *inc* properties were reported for a DNA fragment containing only *ssi* and CR sequences but not for IT1-4 (Rozhon et al., 2011). We can conclude that, regardless of the general structural similarity, pIGMS31, pIGRK, and pHW126 REP modules differ from each other even in the functioning of common core elements. This diversity in the location of *inc* regions, within REP modules of pHW126-like plasmids, clearly indicates that their regulatory networks are organized differently.

The pIGMS31 RepM protein is a multifunctional factor responsible not only for the initiation of replication but also for the up- or down-regulation of the transcriptional activity of REP promoters, including *repM* promoter (Figure 1 and 4). Our EMSA results indicate a direct interaction of recombinant RepM with multiple DNA fragments of the pIGMS31 REP module. They are the IR1 and IR2 sequences (P_MS1_ and P_MS5_ operatory elements) and the IT1-4 iteron-like direct repeats from the replication *origin*. IR and IT sequences share sequence similarity that resembles the organization of Rep protein binding sites in *theta* plasmids (del Solar et al., 1998). In addition, it seems that RepM binds to DNA fragments containing promoters that are activated in the presence of this protein: the CR region comprising the P_MS4_ promoter, and the terminal part of *repM* gene containing the P_MS3_ promoter (Figure 2). Interestingly, no sequence similarity was observed between IR/IT and fragments containing P_MS3_, or P_MS4_. This may suggest that RepM can recognize different types of target DNA sequences. Unfortunately, the unambiguous interpretation of our EMSAs is problematic since we observed unspecific some interaction of purified RepM with the control DNA fragment. On the other hand, a direct interaction on RepM with P_MS3_ and P_MS4_ is the simplest and more likely interpretation for the promoter activation by RepM observed in our *in vivo* experiments.

In addition to the DNA binding activity, we have shown DNA relaxase activity of RepM in *in vitro* nuclease assays. Using 5’ RECE we located DNA nicking within the minimal replication *origin* just upstream the CR sequence (Figure 2).

In our previous work (Wawrzyniak et al., 2019) and under comparative analyses (Supplementary Figure S7) we revealed some similar properties of RepR protein from pIGRK. It acts as a replication initiation factor and also as a transcriptional regulator of pIGRK REP promoters. Rep protein from pHW126 plasmid acts as an autoregulator (Rozhon, 2017). What is more, it showes DNA relaxase activity (Rozhon et al., 2011), while the nuclease activity of RepR are unclear (unpublished results). Sequence similarity analysis showed that pHW126-like Rep proteins, including RepM, are distantly related to the RepL proteins encoded by Staphylococcal pSN2-like plasmids (Rozhon et al., 2010; Rozhon et al., 2011). It was proposed that pSN2 is an archetype of a new RC plasmids family (Khan and Novick, 1982). However, pSN2 and its relatives have been studied to a lesser extent regarding their replication mode (Kwong et al., 2017). Significantly, pHW126-like (and pSN2-like) Rep proteins are deprived of the typical motifs present in RC initiators and the presence of wHTH motifs resemble *theta*-type replication initiators (Kim et al., 2020).

It was initially proposed by Rozhon and co-workers that pHW126-like replicons, similar to pSN2-like plasmids, form a separate family of RC plasmids (Rozhon et al., 2010; Rozhon et al., 2011). However, as mentioned above, the pIGMS31 REP module exhibits some features typical for both RC and *theta* replicons. We decided to determine pIGMS31 replication mode by analyzing the replication intermediates by 2D gel electrophoresis. Surprisingly, we identified replication intermediates forms typical for both *theta* and RC plasmid replication.

Both mutational analysis and 2D gel experiments indicated that DNA sequences located between Sau96 and SpeI sites are required *in cis* for pIGMS31 replication (compare Figure 1A and Figure 3B). In addition, *in vitro* assays positioned RepM nicking site within the DNA region indicated as the RC replication initiation region mapped by 2D gel analysis (compare Figure 2C and Figure 3B). Furthermore, RIs analysis revealed the presence of the other *theta* replication start sites. Multiple replication initiation sites were identified in e.g. R6K plasmid. Its REP module consists of three *origins* activated by the common initiatory factor, π protein. The main *origin* (*oriγ*) can fire independently, while initiation from two other *origins* (*oriα* and *oriβ*) requires the presence *oriγ in cis*-acting (Saxena et al., 2010; Rakowski and Filutowicz, 2013). A similar relationship might link replication origins in pIGMS31. However, our results indicate a much larger number of *theta* initiation sites. it’s more likely that similar to e.g. *E. coli* chromosome, *ori*-independent initiation can occur during pIGMS31 replication (de Massy et al., 194; Asai et al., 1993; reviewed in Ravoitytė and Wellinger, 2017). There are two mechanisms of *ori*-independent initiation involving transcription-driven R-loop formation and/or DNA recombination (Stuckley et al., 2015). Significantly, DNA recombination processes can be involved in pIGMS31 replication initiation since numerous signals corresponding to recombination intermediates were identified in 2D gel experiments (Figure 3). The co-existence of more than one replication mode is unusual for bacterial plasmids, but It is common for other types of prokaryotic replicons – bacterial viruses named bacteriophages. It is believed that this complex replication mode is a consequence of their specific parasitic lifestyle (Weigel and Seitz, 2006). We cannot exclude that pIGMS31 REP and its relatives are satellite phages like e.g. P4 phage, which maintains in lysogeny plasmid-like form and enters into the lytic cycle in the presence of helper P2 phage (Briani et al., 2001). However, it is worth remembering that our knowledge about replication modes of bacterial plasmids comes from testing a few model replicons. So, there may be plasmids that use more complex replication mechanisms, than the simple *theta*, RC, and D-loop modes, as we propose for pIGMS31.

Our second assumption is that closely related plasmid REPs, with significant similarity in replication *origin* as well as initiatory factor structure, can function differently. Our studies on pIGRK and pIGMS31 replication are a great opportunity to track the paths of plasmid REP systems’ evolution and differentiation. Both related plasmids are compatible replicons since both were isolated from the same strain of *K. pneumoniae* (Smorawińska et al., 2012). There are still some questions about REP modules of the pHW126-like plasmids. For example, we unknow if pHW126 REP contains additional promoters, similar to pIGMS31 or pIGRK REP promoters. Complete sets of RNA/protein products expressed by each of pHW126-like REPs should be identified. Nevertheless, based even only on *inc* determinants mapping experiments, it can be argued that the results from studies on one replication system, cannot be easily extrapolated to other closely related systems without experimental confirmation (Figure 4). What is more, knowing the significant difference between pIGMS31, pIGRK, and pHW126 REP modules it is necessary to investigate the replication mode of each plasmid to if determine the members of whole pHW126-like replicons share the same replication mechanism.

Finally, the results presented here provided us with information crucial for future application use of pIGMS31 as a plasmid vector. For example, the nuclease activity of RepM and DNA recombination events associated with plasmid replication should be taken into consideration during the construction of genetic-stable and save transgene carriers based on pIGMS31 (Zaleski et al., 2017). On the other hand, detailed knowledge about the pIGMS31 REP structure and functioning of pIGMS31 REP significantly increases our ability to get rid to level of the negative traits characteristic for this module. It also facilitates make easier to reinforce positive features, such as like copy-up mutants construction, (for theto increase of plasmid yield,) or removal of a fewsome ISS sequences to lower reduce plasmid DNA immunogenicity (Faurez et al., 2010; Zaleski et al., 2017).

## Author Contributions

PW, DB, SB, PH designed the study. PW wrote the draft manuscript. DB, PH supervised the project, discussed, revised, and modified the manuscript to prepare its final version. PW, MD, VCA, ŁS, PK, AG performed the experiments. All authors read and approved the manuscript for publication.

## Acknowledgments

We would like to acknowledge Ł. Dziewit for kindly providing plasmids pcI434SXT and pcI22SXT used as a positive control for *in vivo* detection of protein dimers, and R. Lasek for his help with optimization of the *β*-galactosidase assay.

## Supplementary Material

**Supplementary Figure S1**

Production of TNF-α and INF-γ inducted by DNA of pIGMS31 and pIGRK. **(A)** The number of putative ISS sequences present within pIGMS31 and pIGRK as well as pUC18 genomes. **(B)** Comparison of cytokine production by C57.1 cells transfected with pIGMS31KAN and pIGRKKAN as well as pUC18, K – cells transfected without plasmid (electric shock only).

**Supplementary Figure S2**

SDS-PAGE of recombinant RepM protein purification procedure. Lanes: (1) protein marker, (2) bacterial crude extract from non-inducted culture, (3) bacterial crude extract from IPTG inducted culture, (4) supernatant of centrifuged bacterial lysate, (5) proteins not attached to the Ni-NTA resin, (6, 7) washes, (8) elution.

**Supplementary Figure S3**

The DNA sequence of pIGMS31 plasmid. The minimal replication origin sequence determined by mutational analysis is highlighted in a blue underlined font. Oligonucleotides GB3 and GB9 sequences used for DNA nick site determination are marked by a red font. GTT codon (the equivalent of pIGRK GTG RepR’ start codon) is marked by a blue font. Amino acid residues of *in silico* predicted RepM wHTH motif are marked by a orange font. Codon mutated in pMS-7 is highlighted by a purple underlined font. MobM coding sequence is marked by a grey underlined font. The rest of the markings as in the main text.

**Supplementary Figure S4**

Comparison of rep gene products from pHW126-like plasmids. Rep_pHW126_ – initiatory protein from pHW126, RepR – initiatory protein from pIGRK (sequence of in-frame encoded replication negative regulator RepR’ is highlighted by underlined font), RepM – initiatory protein from pIGMS31, H – alpha helix residue, E – beta-sheet residue, α3 – DNA binding helix, α1,5,6 – dimerization helix, residue marked by the red font is residue mutated in pMS-7.

**Supplementary Figure S5**

Determination of the basic and minimal pIGMS31 replicon. **(A)** Genetic organization of pIGMS31. Black lines represent DNA fragments of pIGMS31 used to determine the minimal replicon. The insertion sites for the kanamycin resistance cassette (KM) and the replication origin of R6K (oriγ) are indicated by black triangles. The ability (+) or inability (-) of the constructed plasmids to replicate in *E. coli* DH5α or *E. coli* DH5α (*repM*) harboring pUC-*repM*_1 is indicated (nd – not determined). **(B)** Comparison of plasmid pMS-3 and its variant pRK-3_2 lacking the *ssi* sequence as well as pMS-5. Electrophoretic separation of purified plasmid DNA on a 1% agarose gel showing the different forms of plasmid DNA. **(C)** Schematic representation of pMS-5γ (*repM* deficient pIGMS31 derivative) and pUC-*repM*_1 (source of RepM) plasmids constructed for *trans*-activation of replication *origin*. In pUC-*repM*_1 ampicillin resistance cassette (AP) and replication *origin* (ori pMB1) were indicated.

**Supplementary Figure S6**

Mapping of pIGMS31 incompatibility determinants. Elements of basic and minimal replicon are indicated: single-strand initiation site (*ssi*), conserved region (CR), iteron-like sequences (IT), and inverted repeats (IR). DNA fragments cloned in the pUC18 vector are marked as solid lines, red lines represent fragments containing incompatibility determinants (*inc1, inc2*). pUC18 derivatives, containing pIGMS31 fragments, were transformed in to DH5α strain harboring pMS-5. Plasmid incompatibility was measured as a loss of kanamycin-resistant clones (harboring pMS-5) in bacterial cultures cultivated for about 80 generations in a non-selective medium.

**Supplementary Figure S7**

Functional analysis of the replication system of plasmid pIGRK **(A)** Mapping of pIGRK incompatibility determinants. Elements of basic and minimal replicon are indicated: single-strand initiation site (*ssi*), conserved region (CR), iteron-like sequences (IT), inverted repeats (IR), and direct repeats (DR). DNA fragments cloned in the pUC18 vector are marked as solid lines, and red lines represent fragments containing incompatibility determinants (*inc1-inc4*). Plasmid incompatibility was measured as a % of Km resistant (harboring pRK-1) clones in bacterial cultures cultivated for about 80 generations in a non-selective medium. **(B)** Identification and analysis of pIGRK REP promoters activity. Promoters identified within the REP module are designed as black arrows. black lines represent DNA fragments of pIGRK and their mutated versions, white arrows set the orientation of DNA fragments cloned in the pRS551 test vector. Black triangles mark the deletion of the HindIII site (frame shist mutation resulting in the lack of RepR), gtg RepR’ internal start codon with a point mutation (result in non-start codon gtc). **(C)** *β*-galactosidase activity of protein extracts from strains carrying pRS551-based constructs imaged in panel (B), reflecting the strength of REP promoters, K – “empty” test vector. **(DE)** REP promoters activity in the presence or absence of RepR, RepR’ or both proteins derived *in trans* from pBAD vector (for more detail see Supplementary Table S1).

**Supplementary Table S1**

Bacterial strains, plasmids, and genetic cassettes used in this study.

**Supplementary Table S2**

Sequences of oligonucleotides used in this study.

