## Supplementary Material for "The unique structure and replication mode of the replication system of *Klebsiella pneumoniae* plasmid pIGMS31"

### 1. Immunogenicity of pIGMS31 and pIGRK derivatives

Mouse mast cells (C57.1) were transfected with DNA of pIGMS31 and pIGRK kanamycin-resistant (KAN) derivatives (260V, 950 µF, ∞ Ω, 30 ms, cuvette 4 mm in DMEM 400 µl of cell suspensions, 25 µg plasmid DNA). After transfection cells were incubated in a cDMEM medium. Measurement of secreted cytokines was performed after 2 days using Mouse Th1/Th2/Th17 Cytokine Kit (Becton, Dickinson and Company, catalog no. 560485). The Number of C57.1 cells was estimated at $2x10^6$/sample.

(A)

| ISS: | pIGMS31 | pIGRK | pUC18 |
|---|---|---|---|
| RRCGYY | 0 | 2 | 14 |
| GTCGYY | 2 | 0 | 6 |
| RRCGTT | 0 | 2 | 9 |

(B)

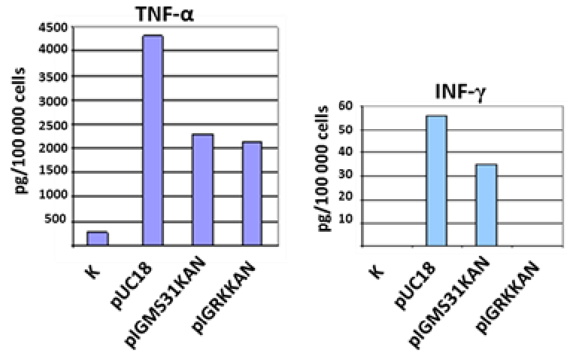


**Supplementary Figure S1**. Production of TNF-α and INF-γ inducted by DNA of pIGMS31 and pIGRK. (A) The number of putative ISS sequences present within pIGMS31 and pIGRK as well as pUC18 genomes. (B) Comparison of cytokine production by C57.1 cells transfected with pIGMS31KAN and pIGRKKAN as well as pUC18, K – cells transfected without plasmid (electric shock only).

**2. RepM(6His) purification**

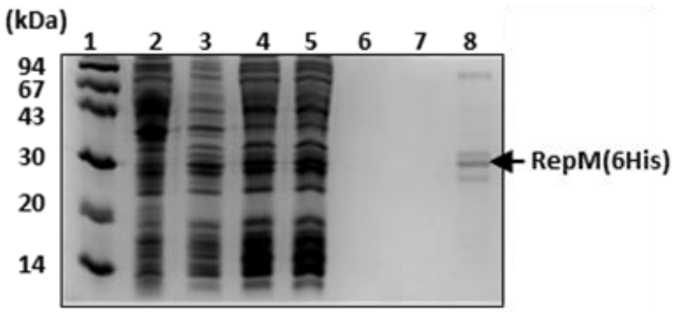


**Supplementary Figure S2.** SDS-PAGE of recombinant RepM protein purification procedure. Lanes: (1) protein marker, (2) bacterial crude extract from non-inducted culture, (3) bacterial crude extract from IPTG inducted culture, (4) supernatant of centrifuged bacterial lysate, (5) proteins not attached to the Ni-NTA resin, (6, 7) washes, (8) elution.

### 3. Genetic organization of pIGMS31 plasmid

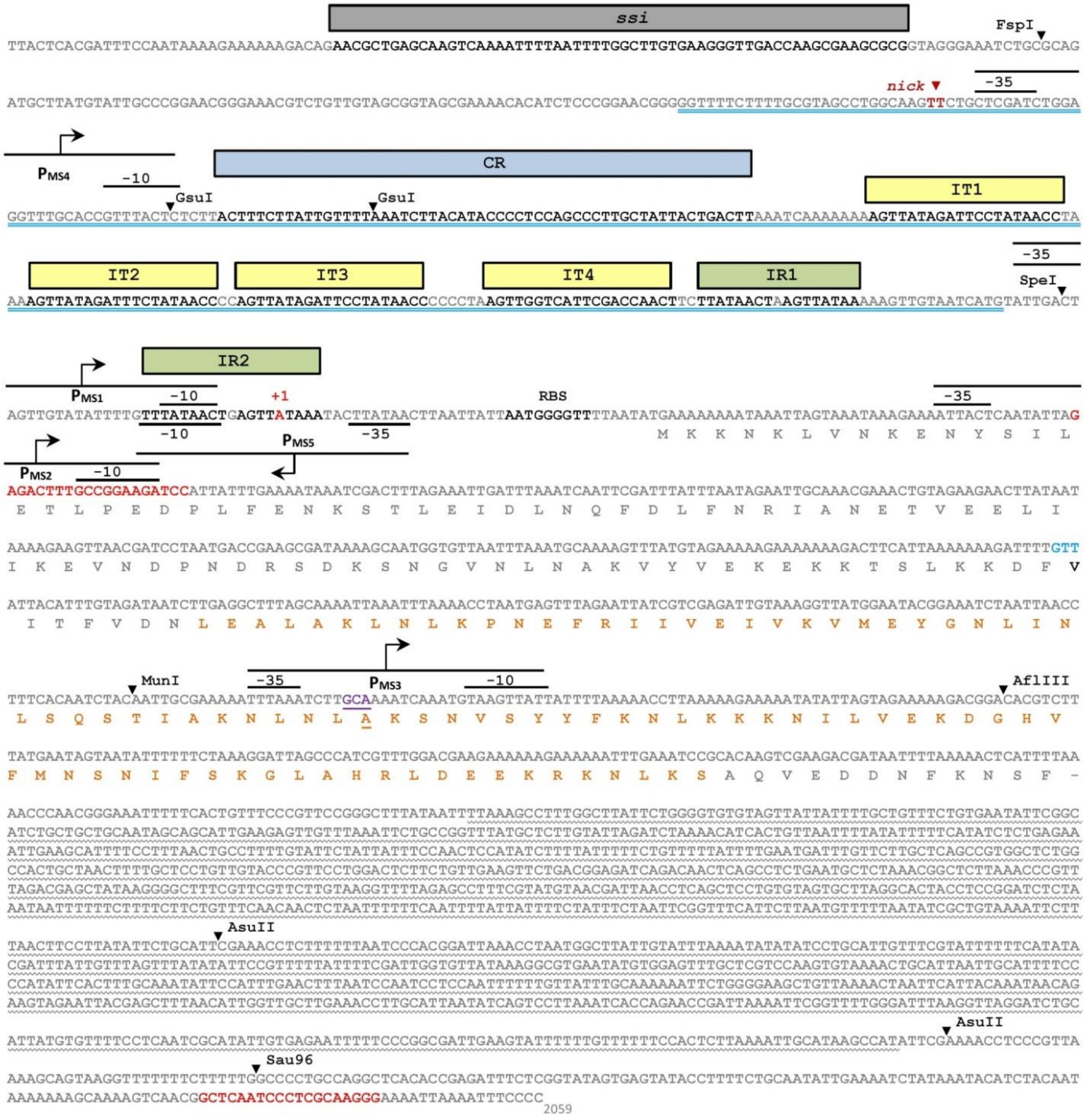


**Supplementary Figure S3.** The DNA sequence of pIGMS31 plasmid. The minimal replication *origin* sequence determined by mutational analysis is highlighted in a blue underlined font. Hybridization sites of GB3 and GB9 oligonucleotides used for 5’ RACE DNA nick site determination are marked by a red font. GTT codon (the equivalent of pIGRK GTG RepR’ start codon) is marked by a blue font. Amino acid residues of *in silico* predicted RepM wHTH motif are marked by a orange font. Codon mutated in pMS-7 is highlighted by a purple underlined font. MobM coding sequence is marked by a grey underlined font. The rest of the markings as in the main text.

### 4. Comparison of *rep* gene products from pHW126-like plasmids

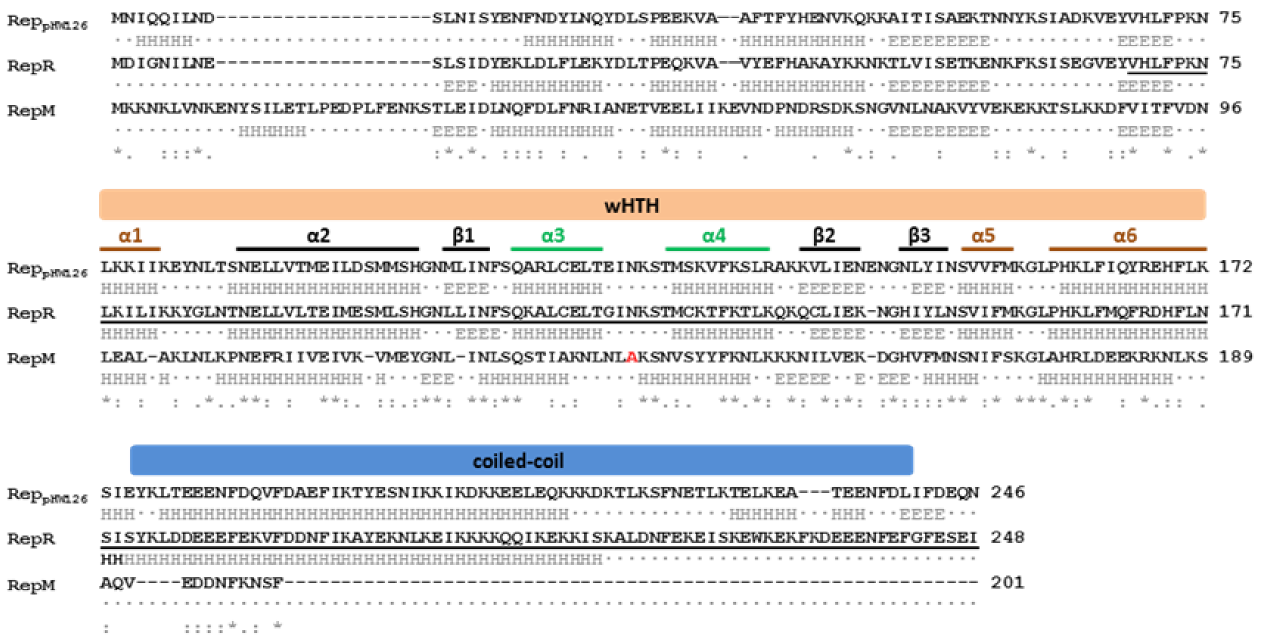


**Supplementary Figure S4.** Comparison of rep gene products from pHW126-like plasmids. $Rep_{pHW126}$ – initiatory protein from pHW126, RepR – initiatory protein from pIGRK (sequence of in-frame encoded replication negative regulator RepR' is highlighted by underlined font), RepM – initiatory protein from pIGMS31, H –alpha helix residue, E – beta-sheet residue, α3 – DNA binding helix, α1,5,6 – dimerization helix, residue marked by a red font is residue mutated in pMS-7.

### 5. Detailed imaging of molecular analysis of pIGMS31 replication system elements

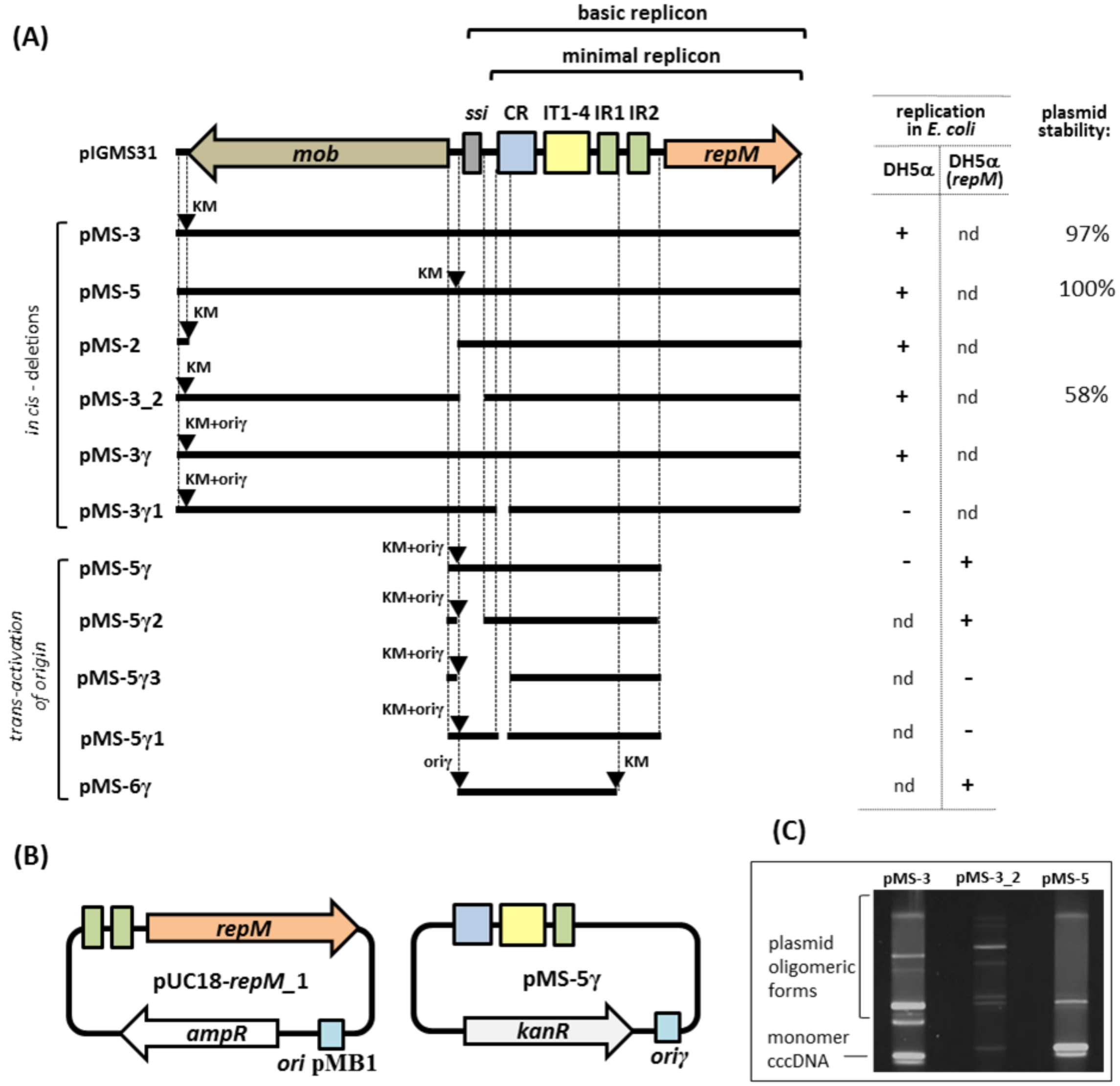


**Supplementary Figure S5.** Determination of the basic and minimal pIGMS31 replicon. **(A)** Genetic organization of pIGMS31. Black lines represent DNA fragments of pIGMS31 used to determine the minimal replicon. The insertion sites for the kanamycin resistance cassette (KM) and the replication origin of R6K *(oriγ)* are indicated by black triangles. The ability (+) or inability (-) of the constructed plasmids to replicate in E. coli DH5α or E. coli DH5α (*repM*) harboring pUC-*repM*_1 is indicated (nd – not determined). **(B)** Comparison of plasmid pMS-3 and its variant pRK-3_2 lacking the ssi sequence as well as pMS-5. Electrophoretic separation of purified plasmid DNA on a 1% agarose gel showing the different forms of plasmid DNA. **(C)** Schematic representation of pMS-5γ (*repM* deficient pIGMS31 derivative) and pUC-*repM*_1 (source of RepM) plasmids constructed for trans-activation of replication origin. In pUC-*repM*_1 ampicillin resistance cassette (AP) and replication *origin* (*ori* pMB1) were indicated.

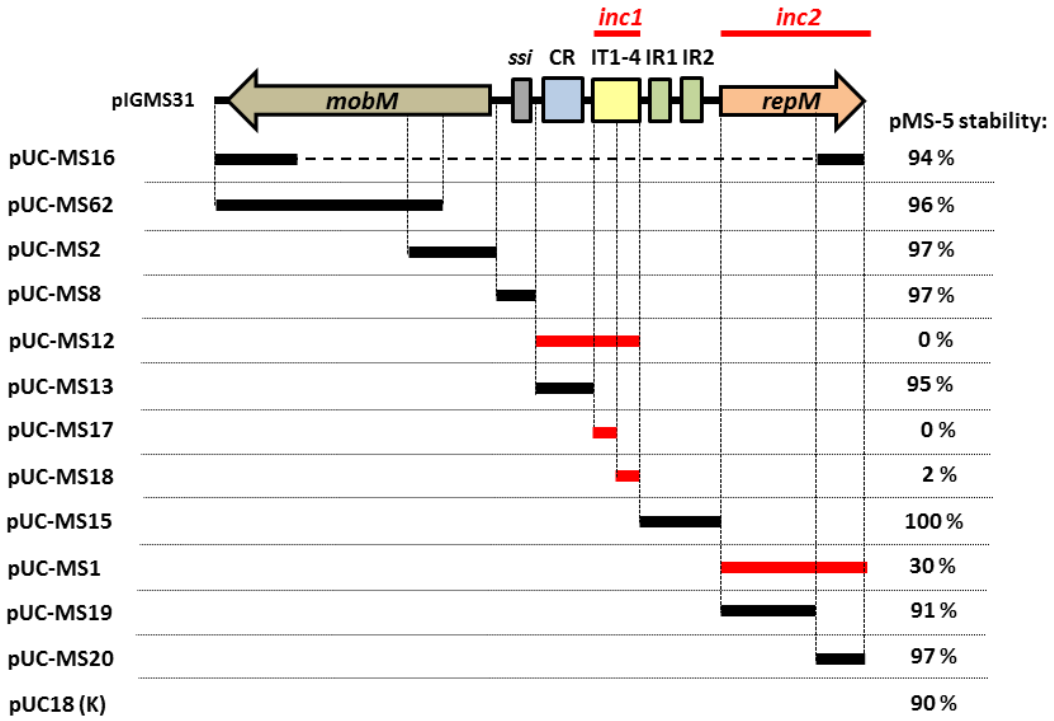


**Supplementary Figure S6.** Mapping of pIGMS31 incompatibility determinants. Elements of basic and minimal replicon are indicated: single-strand initiation site (*ssi*), conserved region (CR), iteron-like sequences (IT), and inverted repeats (IR). DNA fragments cloned in the pUC18 vector are marked as solid lines, red lines represent fragments containing incompatibility determinants (inc1, inc2). pUC18 derivatives, containing pIGMS31 fragments, were transformed in to DH5α strain harboring pMS-5. Plasmid incompatibility was measured as a loss of kanamycin-resistant clones (harboring pMS-5) in bacterial cultures cultivated for about 80 generations in a non-selective medium.

### 6. Molecular analysis of pIGRK replication system elements

Incompatibility determinants analyses, as well as promoter identification, were performed according to the procedures implemented for pIGMS31 studies and described in the main text of this paper as well as procedures described in our last publication concerning research on pIGRK plasmid (Wawrzyniak et al., 2019)

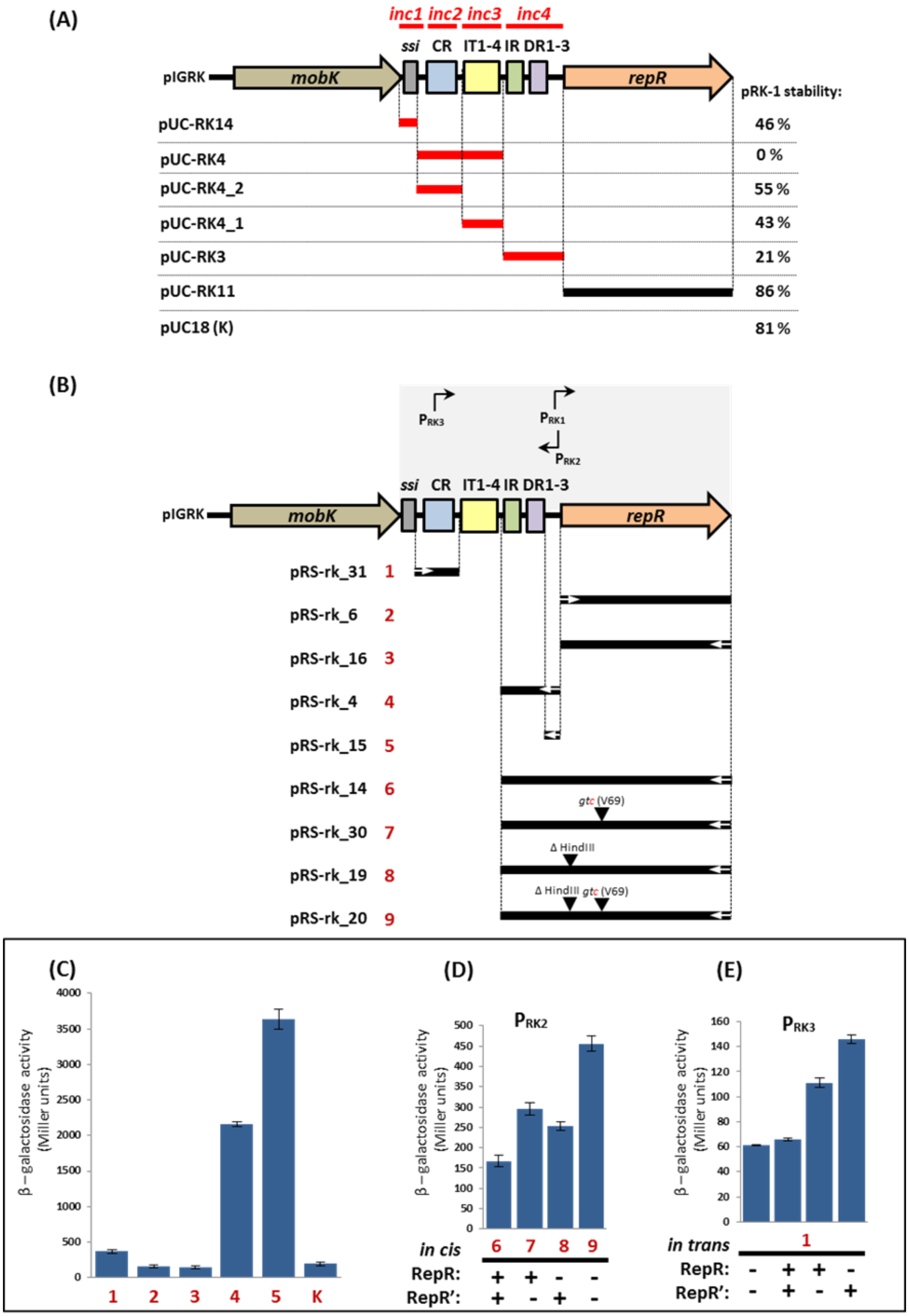


**Supplementary Figure S4**. Functional analysis of the replication system of plasmid pIGRK **(A)** Mapping of pIGRK incompatibility determinants. Elements of basic and minimal replicon are indicated: single-strand initiation site (*ssi*), conserved region (CR), iteron-like sequences (IT), inverted repeats (IR), and direct repeats (DR). DNA fragments cloned in the pUC18 vector are marked as solid lines, and red lines represent fragments containing incompatibility determinants (inc1-inc4). Plasmid incompatibility was measured as a % of Km resistant (harboring pRK-1) clones in bacterial cultures cultivated for about 80 generations in a non-selective medium. **(B)** Identification and analysis of pIGRK REP promoters activity. Promoters identified within the REP module are designed as black arrows.  black lines represent DNA fragments of pIGRK and their mutated versions, white arrows set the orientation of DNA fragments cloned in the pRS551 test vector. Black triangles mark the deletion of the HindIII site (frame shist mutation resulting in the lack of RepR), gtg RepR' internal start codon with a point mutation (result in non-start codon gtc). **(C)** *β*-galactosidase activity of protein extracts from strains carrying pRS551-based constructs imaged in panel (B), reflecting the strength of REP promoters, K – "empty" test vector. **(D, E)** REP promoters activity in the presence or absence of RepR, RepR' or both proteins derived in trans from pBAD vector (for more detail see Supplementary Table S1).

**Supplementary Table S1.** Bacterial strains, plasmids, and genetic cassettes used in this study.

| **Bacterial strain** | **Description** | |
|---|---|---|
| ***E. coli* DH5α** | $F^-$ *endA1 glnV44 thi-1 recA1 relA1 gyrA96 deoR nupG* Φ80d *lacZ*ΔM15 Δ(*lacZYA-argF*)*U169*, *hsdR17* ($r_K^-$ $m_K^+$), λ | Invitrogen |
| ***E. coli* DH5αΔ*lac*** | *deoR thi1 relA1 supE44 endA1 gyrA96 recA1 hsdR17 Δ(argF lac)U169* $Nal^r$ | M. Yarmolinski* |
| ***E. coli* R721** | *supE thy* Δ(*lac–proAB*) F′ [*proAB*$^+$ *lacI*$^q$ *lacZ*ΔM15] 71/18 *glpT*::O-$P_{434/P22}$ *lacZ* | [1] |
| ***E. coli* BL21(DE3)** | $F^-$ *ompT gal dcm lon* $hsdS_B$ ($r_B^-$ $m_B^-$) λ(*DE3*) *pLysS*($Cm^R$) | Invitrogen |
| ***E. coli* MC1061** | *$F^-$ Δ(ara-leu)7697 $[araD139]_{B/r}$ Δ(codB-lacI)3 galK16 galE15 $λ^-$ $e14^-$ mcrA0 relA1 rpsL150(strR) spoT1 mcrB1 hsdR2($r^-m^+$)* | [2] |
| ***E. coli* DH5αλ*pir*** | *sup E44* Δ*lacU169*(Φ*lacZ*ΔM15) *recA1 endA1 hsdR17 thi-1 gyrA96 relA1 λpir* phage lysogen | [3] |
| **Plasmid** | **Description** | |
| **pIGRK** | cryptic plasmid isolated from *Klebsiella pneumoniae* 287-w | [4] |
| **pIGMS31** | cryptic plasmid isolated from *Klebsiella pneumoniae* 287-w | [4] |
| **pUC18** | $Ap^r$, *ori* pMB1, *lacZ* (M13mp18/19), cloning vector | Qiagen |
| **pABB19** | $Ap^r$; *ori* pMB1, cloning vector with T*pro*/T*lyz* (T) transcriptional terminator from bacteriophage P1, used for cloning of DNA fragments with T sequence | [5] |
| **pET24b+** | $Km^r$, *ori* pBR, *ori* F1, plasmid vector for 6His tagged proteins overexpression, used for Rep proteins identification and purification | Invitrogen |
| **$pcI_{434}$** | $Km^r$, *ori* p15A, pACYC177 derivative coding *N*-terminal part of $P_{434}$ phage repressor, used in bacterial two hybrid system | [1] |
| **$pcI_{22}$** | $Ap^r$, *ori* ColE1, pC132 derivative coding N-terminal part of $P_{22}$ phage repressor, used in bacterial two hybrid system | [1] |
| **$pcI_{434}$SXT** | $pcI_{434}$ derivative coding protein from SXT N15 prophage | [6] |

| | | |
|---|---|---|
| **pcI$_{22}$SXT** | pcl$_{22}$ derivative coding protein from SXT N15 prophage | [6] |
| **pRS551** | Ap$^r$, Km$^r$ , *ori* pBR, test vector coding promoter less *lacZ* reporter gene, used for determination of $P_{repR}$ promoter activity | [7] |
| **pDrive** | Ap$^r$, Km$^r$, *ori* pUC coding vector | Qiagen |
| **pDrivedxs** | pDrive with cloned fragment of *dxs E. coli* gene | [8] |
| **pBAD33** | Cm$^r$, *ori* p15A, PBAD-*araC*, expression vector used for arabinose inducted Rep proteins *in trans* delivery | [9] |
| **pDS132** | Cm$^r$ *ori* R6K, *mobRP4*, *sacB*, used for oriγ cassette construction | [10] |
| **pMS-3** | pMS-2 with SpeI-XhoI replaced by SpeI-XhoI fragment from pMS-1 | this study |
| **pMS-2** | pIGMS31 DNA was PCR amplified using oligo 1 and 2 (Table S2), BamHI digested and ligated with BglII digested KM cassette (700_1867del, 699_1868insKM) | this study |
| **pMS-1** | pIGMS31 DNA was PCR amplified using oligo 3 and 4 (Table S2), BamHI digested and ligated with BglII digested KM cassette (684_689del, 683_690insKM) | this study |
| **pMS-5** | pIGMS31DNA was PCR amplified using oligo 5 and 6 (Table S2), ), BamHI digested and ligated with BglII digested KM cassette (2011_2012insKM) | this study |
| **pMS-3_2** | pMS-3 DNA was PCR amplified using oligo 9 and 10 (Table S2),and self-ligated (2082_2154del) | this study |
| **pMS-3γ** | pMS-3 with oriγ cassette inserted in SmaI site | this study |
| **pMS-3γ1** | pMS-3γ was deprived of GsuI fragment, sticky ends were completely filled-in using Klenow fragment of *E. coli* DNA polymerase and then self-ligated (2299_2322del) | this study |
| **pMS-5γ** | pMS-5 with AsuII-SpeI fragment replaced by oriγ cassette (2511_1873del) | this study |
| **pMS-5γ1** | pMS-5γ was deprived of GsuI fragment, sticky ends were completely filled-in using Klenow fragment of *E. coli* DNA polymerase and then self-ligated (2299_2322del) | this study |
| **pMS-5γ2** | pMS-5γ was deprived of EcoRI-FspI fragment, sticky ends were completely filled-in using Klenow fragment of *E. coli* DNA polymerase and then self-ligated (2013_2167del) | this study |
| **pMS-5γ3** | pMS-5γ was deprived of EcoRI-GsuI fragment, sticky ends were completely filled-in using Klenow fragment of *E. coli* DNA polymerase and then self-ligated (2013_2322del) | this study |
| **pMS-6γ** | Plasmid containing minimal replication *origin* of pIGMS31. Fragment of pMS-3γ, containing pIGMS31 sequence (2242-2509 bp) and oriγ was PCR amplified using oligo 8 and 13 (Table S2) and ligated with KM casette | this study |
| **pMS-7** | pMS-5 derivative, spontanous mutant with substitution of single nucleotide 471G>C in repM ORF (result in RepM 140A>P mutation) | this study |
| **pUC-*repM*_1** | pUC18 with PCR amplified fragment of pIGMS31 (2465-662 bp) cloned in EcoRI and BamHi sites using oligo 14 and 15 (Table S2) | this study |
| **pUC-*repM*_2** | pRS-ms_6 with *repM* start codon replaced by a stop codon, mutation introduced by site-directed mutagenesis and PCR using oligo 22 and 23 (Table S2) | this study |
| **pET-*repM*$_{6H}$** | pET24a+ with PCR amplified *repM* gene (using 16 and 17 oligo, Table S2) (31-662 bp pIGMS31) cloned in BamHI and XhoI sites | this study |
| **pBAD-*repM*** | pBAD33 with insertion of PCR amplified *repM* gene (using 18 and 19 oligo, Table S2 31-662 bp pIGRK) in KpnI and HindIII sites | this study |
| **pRS-ms_1** | pRS551 with pIGMS31 fragment (2486-56 bp) PCR amplified using oligo 20 and 21, Table S2) cloned using EcoRI and BamHI | this study |
| **pRS-ms_2** | pRS551 with pIGMS31 fragment (2465-56 bp) PCR amplified using oligo 14 and 21, Table S2) cloned using EcoRI and BamHI | this study |
| **pRS-ms_3** | Spontaneous mutant of pRS-ms_2 with IS*1* inserted in cloned fragment of pIGMS31 (10_11insIS*1*) | this study |
| **pRS-ms _4** | pABB19PMS1 fragment was PCR amplified and cloned in EcoRI and BamHI pRS551 sites using EcoRI nd BglII. | this study |
| **pRS-ms _6** | pRS551 with pIGMS31 fragment (2465-662 bp) PCR amplified using oligo 14 and 15 Table S2) cloned using EcoRI andi BamHI | this study |
| **pRS-ms_61** | pRS-ms_6 with *repM* start codon replaced by a stop codon, mutation introduced by site-directed mutagenesis and PCR using oligo 22 and 23 (Table S2) | this study |
| **pRS-ms_62** | pRS551 with pIGMS31 fragment (2486-662 bp) PCR amplified using oligo 20 and 15, Table S2) cloned using EcoRI and BamHI | this study |
| **pRS-ms_63** | pRS551 with pIGMS31 fragment (2486-662 bp) PCR amplified using oligo 24 and 25, Table S2) cloned using EcoRI and BamHI | this study |
| **pRS-ms_17** | pRS551 with pIGMS31 fragment (2486-56 bp) PCR amplified using oligo 21 and 26, Table S2) cloned using BamHI | this study |
| **pRS-ms_20** | pUC-MS1 was digested by HindIII (Pol Klenov fragment blunted) and EcoRI, obtained fragment (containing pIGMS31 31-622 bp sequence) and cloned in BamHI (Pol Klenov fragment blunted) and EcoRI | this study |

| | | |
|---|---|---|
| **pRS-ms_21** | pRS551 with pIGMS31 fragment (31-662 bp) PCR amplified using oligo 13 and 29, Table S2) cloned using BamHI and EcoRI sites | this study |
| **pRS-ms_23** | EcoRI-HpaI restriction fragment of pRS-ms_20 (31-221 bp pIGMS31) was cloned in EcoRI and BamHI (Pol Klenov fragment blunted) of pRS551 | this study |
| **pRS-ms_26** | pRS551 with pIGMS31 fragment (2242-2366 bp) PCR amplified using oligo 21 and 26, Table S2) cloned using BamHI | this study |
| **$pcI_{434}M$** | $pcI_{434}$ was digested with SalI and BamHI and ligated with SalI and BamHI digested PCR product (54-662 bp pIGMS31) obtained using oligo 30 and 31 (Table S2) | this study |
| **$pcI_{22}M$** | $pcI_{22}$ was digested with SalI and BamHI and ligated with SalI and BamHI digested PCR product (54-662 bp pIGMS31) obtained using oligo 30 and 31 (Table S2) | this study |
| **pUC-MS_1** | pUC18 with insertion of PCR amplified *repM* gene (using 18 and 19 oligo, Table S2 31-662 bp pIGRK) in KpnI and HindIII sites | this study |
| **pUC-MS_2** | pUC18 digested with SmaI and ligated with pMS-3 AsuII fragment (ends blunted with Pol Klenov fragment) (1354-1877 bp pIGMS31) | this study |
| **pUC-MS_12** | pUC18 with pIGMS31 fragment (2242-2498 bp) PCR amplified using oligo 13 and 32, Table S2) cloned using BamHI and EcoRI | this study |
| **pUC-MS_13** | pUC18 with pIGMS31 fragment (2242-2365 bp) PCR amplified using oligo 13 and 33, Table S2) cloned using BamHI and EcoRI | this study |
| **pUC-MS_6** | pUC18 digested with BamHI and SmaI and ligated with pMS-4 BamHI-AsuII fragment (675-1787 bp pIGMS31) | this study |
| **pUC-MS_62** | pUC18 digested with HindIII and SmaI and ligated with pUC-MS_6 XmnI-HindIII fragment (222-1787 bp pIGMS31) | this study |
| **pUC-MS_16** | pUC18 digested with SmaI and ligated with pMS-5 BamHI-MunI fragment (ends blunted with Pol Klenov fragment) (449-836 bp pIGMS31) | this study |
| **pUC-MS_14** | pUC18 with pIGMS31 fragment (2486-56 bp) PCR amplified using oligo 20 and 21, Table S2) cloned using EcoRI and BamHI | this study |
| **pUC-MS_15** | pUC18 with pIGMS31 fragment (2465-56 bp) PCR amplified using oligo 14 and 21, Table S2) cloned using EcoRI and BamHI | this study |
| **pUC-MS_17** | pUC18 digested with SmaI and ligated with annealed oligo 34 and 35 (Table S2) (2366-2419 bp pIGMS31) | this study |
| **pUC-MS_18** | pUC18 digested with SmaI and ligated with annealed oligo 36 and 37 (Table S2) (2419-2467 bp pIGMS31) | this study |
| **pUC-MS_19** | pUC-MS1 without HindIII-MunI fragment, ends blunted with Pol Klenov fragment were self-ligated (449-662 bp pIGMS31) | this study |
| **pUC-MS_20** | pUC-MS1 without EcoRI-MunI fragment (31_448del bp pIGMS31), ends blunted with Pol Klenov fragment were self-ligated | this study |
| **pUC-RK14** | pIGRK DNA was digested with HphI, the obtained fragment was cloned in the pUC18 SmaI site (sticky ends of the cloned fragment were blunted with Pol Klenov fragment) | this study |
| **pUC-RK4** | pUC18HpaI with DNA fragment of pRK3_2γ PCR amplified using oligo 38 and 39 (Table S2) (2049-2348 bp pIGRK) | [11] |
| **pUC-RK4AsuII** | pUC-RK4 with AsuII site (2257A>C, 2258A>G) introduced by PCR using oligo 40 and 41 (Table S2) | [11] |
| **pUC-RK4_1** | pUC-RK4AsuII was digested with AsuII and EcoRI, sticky ends of the plasmid vector were blunted with Pol Klenov fragment and self-ligated (2259_2348del) | [11] |
| **pUC-RK4_2** | pUC-RK4AsuII was digested with HindIII and EcoRI, sticky ends of the plasmid vector were blunted with Pol Klenov fragment and self-ligated (2049_2257del) | [11] |
| **pUC-RK3** | pUC18 with pIGRK fragment (10-136 bp) PCR amplified using oligo 42 and 43 (Table S2) cloned using EcoRI | this study |
| **pUC-RK11** | pUC18 with pIGRK fragment (125-888 bp) PCR amplified using oligo 44 and 45 (Table S2) cloned using EcoRI and BamHI | this study |
| **pRS-rk_31** | the pUC-RK4_1 DNA fragment was PCR amplified using oligo 46 and 47 (Table S2) and cloned into pRS551 BamHI (sticky end of the plasmid vector was blunted with Pol Klenov fragment) and EcoRI sites | this study |
| **pRS-rk_6** | pRS551 with pIGRK fragment (61-888 bp) PCR amplified using oligo 48 and 49 (Table S2) and cloned using EcoRI and BamHI | [11] |
| **pRS-rk_16** | pRS-rk_14 vector with deletion of BamHI-BseRI restriction fragment (1_140del), sticky ends of the plasmid vector were blunted with Pol Klenov fragment and self-ligated | this study |
| **pRS-rk_4** | pRS551 with pIGRK fragment (1-141 bp) PCR amplified using oligo 42 and 43 (Table S2) and cloned using EcoRI | this study |

| | | |
|---|---|---|
| **pRS-rk_8** | pRS551 with cloned EcoRI-BamHI fragment of pUC-*repR*_2 [11] | this study |
| **pRS-rk_15** | pRS-rk_8 lacking EcoRI-SpeI fragment (1_77del), sticky ends of the plasmid vector were blunted with Pol Klenov fragment self-ligated | this study |
| **pRS-rk_14** | pRS551 with pIGRK fragment (61-888 bp) PCR amplified using oligo 48 and 49 (Table 2) and cloned using EcoRI and BamHI | this study |
| **pRS-rk_30** | pRS551 with pUC-*repR*_2 [11] fragment cloned in EcoRI-BamHI sites, insert was PCR amplified using oligo 50 and 51 (Table S2) | this study |
| **pRS-rk_19** | pRS551 with pUC-*repR*_8 [11] fragment cloned in EcoRI-BamHI sites, insert was PCR amplified using oligo 50 and 51 (Table S2) | this study |
| **pRS-rk_20** | pRS551 with pUC-*repR*_3 [11] fragment cloned in EcoRI-BamHI sites, insert was PCR amplified using oligo 50 and 51 (Table S2) | this study |
| **pBAD-*repR*** | pBAD33$_{\Delta H}$ with *repR* gene cloned in XbaI and PstI sites, insert was PCR amplified using oligo 48 and 49 (Table S2) | [11] |
| **pBAD-*repR*$_{V69V}$** | pBAD-*repR* with V69V mutation (343G>C) | [11] |
| **pBAD-*repR*$_{\Delta H}$** | pBAD-*repR* digested with HindIII, sticky ends of the plasmid vector were blunted with Pol Klenov fragment and self-ligated | [11] |
| **pUC18HpaI** | pUC18 with additional HpaI restriction site introduced by PCR with oligo 52 and 53 (Table S2) (419C>T, 420G>A, ΔAccI, ΔHincII, ΔSalI) | [11] |
| **pBAD33$_{\Delta H}$** | pBAD33 was digested with HindIII, sticky ends were filled in using Klenow fragment of *E. coli* DNA polymerase, and then self-ligated | [11] |
| **pABB19*kan*** | pABB19 was digested with BamHI and ligated with BamHI digested PCR product (EZ-Tn5™ <KAN-2> transposon 20-1051 bp fragment, amplified using oligo 54 and 55, Table S2) | [11] |
| **pABB19P$_{MS1}$** | pABB19 digested with EcoRI, BamHI and ligated with EcoRI, BamHI digested PCR product (2465-56 bp pIGMS31 fragment, amplified using oligo 14 and 21 (Table S2) | this study |
| **Cassette** | **Description** | |
| **KM cassette** | pABB19*kan* fragment containing kanamycin resistance *kan* gene and transcriptional terminator (T) PCR amplified using oligo 56 and 39 (Table S2) containing BglII sites | [11] |
| **oriγ cassette** | pDS132 fragment (1-392 bp) PCR amplified using oligo 7 and 8 (Table S2) containing SpeI and AsuII sites | [11] |

The introduced mutations are described according to the following scheme: (i) deletions: 000_000del (nucleotide position of the first deleted pair of bases_nucleotide position of the last deleted pair of bases, deletion), (ii) insertions: 000_000insXYZ (nucleotide position of the first pair of bases above the insertion_position of the first pair of bases below the insertion, ins - insertion; XYZ - the name of the inserted element), (iii) nucleotide substitutions: 000X> Y (000X - position in the sequence and nucleotide occurring in the sequence originally, Y - the nucleotide introduced in its place), (iv) amino acid substitutions: X00Y (X - original amino acid, 00 - position in the sequence, Y - introduced amino acid). Sequence coordinates, unless otherwise stated, pIGMS31 (GenBank: AY543072.1), *unpublished data.

**Supplementary Table S2.** Sequences of oligonucleotides used in this study.

| | Name | Oligonucleotide sequence (5′→3′) |
|---|---|---|
| **1** | MSB_F | GA**GGATCC**GCCATATTCGAAAACCTCCC |
| **2** | MSB_R | GA**GGATCC**AGCCCGGAACGGGAAAC |
| **3** | MSZ_F2 | GA**GGATCC**GTTCCGGGCTTTATAATTTTAAAGC |
| **4** | MSZ_R2 | GA**GGATCC**CAGTGAAAAATTTCCCGTTGGG |
| **5** | MSWT3F | GAT**GGATCC**CAAAAGTCAACGGCTCAATC |
| **6** | MSWT3R | GAT**GGATCC**GCTTTTTTTATTGTAGATGTATTTATAG |
| **7** | r6kSpeIF | GAT**ACTAGT**CCATGTCAGCCGTTAAGTGTTCC |
| **8** | r6kAsuR | GAT**TTCGAA**GATCTGAAGATCAGCAGTTCAACCTG |
| **9** | MSssiR | GGT**AGATCT**GTTTTATTGGAAATCGTGAGTAAGG |
| **10** | MSssiF | GAT**AGATCT**AGGGAAATCTGCGCAGATGC |
| **11** | MVGTAF | ATCTATTTCCGAAGGTGTTGAATACGT**c**CATTTATTCCC |

| | Name | Oligonucleotide sequence (5′→3′) |
|---|---|---|
| **12** | MVGTAR | GGGAATAAATG**g**ACGTATTCAACACCTTCGGAAATAGAT |
| **13** | MSNIKFE | GTC**GAATTC**GGTTTTCTTTTGCGTAGCCTGG |
| **14** | PMSUPEF | GAT**GAATTC**TTCTTATAACTAAGTTATAAAAAGTTG |
| **15** | PmsrepMR | TAA**GGATCC**GTTTTAAAATGAGTTTTAAAATTATCG |
| **16** | REPM_F2 | GA**GGATCC**TTAATTATTAATGGGGTTTTAATATG |
| **17** | REPMPETR | GA**CTCGAG**AATGAGTTTTTAAAATTATCGTCT |
| **18** | RBSrepMF | ATAT**GGTACC**TTAATTATTAATGGGGTTTTAATATG |
| **19** | RBSrepMR | TGGC**AAGCTT**GTTTTAAAATGAGTTTTAAAATTATCG |
| **20** | PmsrepMF | GTA**GAATTC**AAGTTGTAATCATGTATTGACTAG |
| **21** | PmsBamHR | GAT**GGATCC**ATATTAAAACCCCATTAATAATTAAG |
| **22** | MutTAGF | CTTTATTTACTAATTTATTTTTTTTCT**a**ATTAAAACC |
| **23** | MuTAGR | GGTTTTAAT**t**AGAAAAAAAATAAATTAGATAAATAG |
| **24** | PmsrepMF2 | GTA**GGATCC**AAGTTGTAATCATGTATTGACTAG |
| **25** | PmsrepMR2 | TAA**GAATTC**GTTTTAAAATGAGTTTTAAAATTATCG |
| **26** | PMSUPBF | CAATAGATTAAGTTGCAAAAAGTAGAAAAAGTTG |
| **27** | EPmsrepR | GTA**GAATTC**GTTTTAAAATGAGTTTTTAAAATTATCG |
| **28** | RBSrepMBF | ATAT**GGATCC**TTAATTATTAATGGGGTTTTAATATG |
| **29** | PMSUPEF | GTA**GAATTC**TTCTTATAACTAAGTTATAAAAAGTTG |
| **30** | RepMFSalI | GC**GTCGAC**CATGAAAAAAAATAAATTAGTAAATAAAGAAAATTAC |
| **31** | RepMRBam | TAA**GGATCC**GTTTTAAAATGAGTTTTTAAAATTATCGTC |
| **32** | MSDRRB0 | GCG**GGATCC**ATGATTACAACTTTTTATAACTTAGTTATAAGAAG |
| **33** | MSNIKRB | GAT**GGATCC**GATTTAAGTCAGTAATAGCAAGGGC |
| **34** | MSTI12F | CAAAAAAAAGTTATAGATTCCTATAACCTAAAAGTTATAGATTTCTATAACCC |
| **35** | MSIT12R | GGGTTATAGAAATCTATAACTTTTAGGTTATAGGAATCTATAACTTTTTTTTG |
| **36** | MSIT34F | CAGTTATAGATTCCTATAACCCCCCTAAGTTGGTCATTCGACCAACTTC |
| **37** | MSIT34R | GAAGTTGGTCGAATGACCAACTTAGGGGGGTTATAGGAATCTATAACTG |
| **38** | DsoRK_R | CTAGAGTTGTCGGATTTGACAACCTC |
| **39** | Pabbkasf | GTA**AGATCT**CGACGGCCAGTGAATTCGAGCTC |
| **40** | RKASUIIF | CTCTAACCATTGATATTACTGGATTTTT**cg**AAAAGGCAGTTGTC |
| **41** | RKASUIIR | GACAACTGCCTTTT**cg**AAAAATCCAGTAATATCAATGGTTAGAG |
| **42** | TNREPL | AA**GAATTC**AGACTCTAGCCAGTTTC |
| **43** | TNREPR | CT**GAATTC**TCCATAGAAACCTCCTC |
| **44** | BEcrepRF | GCT**GAATTC**GAGGAGGTTTCTATGGATATTGG |
| **45** | PRSrepRR | GAT**GGATCC**CGGTTTTATATTTCCGATTCAAAACC |
| **46** | RKssiF2 | GTA**GAATTC**AGCAAAATCCTCCATAGCGAAG |
| **47** | M13pUCf | CCAGTCACGACGTTGTAAAACG |
| **48** | BEcrepRF | GCT**GAATTC**GAGGAGGTTTCTATGGATATTGG |
| **49** | PRSrepRR | GAT**GGATCC**CGGTTTTATATTTCCGATTCAAAACC |
| **50** | TNREPBf | AA**GGATCC**AGACTCTAGCCAGTTTC |
| **51** | repRRECO | GCG**GAATTC**CGGTTTTATATTTCCGATTCAAAACC |
| **52** | pucHpaIR | GATCCTCTAGAGT**ta**ACCTGCAGGCATGCAAGCTTG |
| **53** | pucHpaIF | CAAGCTTGCATGCCTGCAGGT**ta**ACTCTAGAGGATC |
| **54** | KANBAM_F | GAG**GGATCC**CAACCATCATCGATGAATTG |
| **55** | KANBAM_R | GAG**GGATCC**CTTCAACTCAGCAAAAGTTC |
| **56** | Pabbkasr | GTA**AGATCT**GTTCCTGGCCTTTCCATGGATAATAGTTAACG |
| **57** | r6kSpeIF | GAT**ACTAGT**CCATGTCAGCCGTTAAGTGTTCC |
| **58** | r6kAsuR | GAT**TTCGAA**GATCTGAAGATCAGCAGTTCAACCTG |
| **59** | M13pUCr | AGCGGATAACAATTTCACACAGG |
| **60** | M13pUCfFAM | FAM-CCAGTCACGACGTTGTAAAACG |
| **61** | GB3 | GCTCAATCCCTCGCAAGGG |
| **62** | GB9 | CCCTTGCGAGGGATTGAGC |
| **63** | SP2RACEMS | CTTCGTCCAAACGATGGGCTAATCCTTTAG |
| **64** | Oligo d(G) | GACCACGCGTATCGATGTCGACGGGGGGGGGGGGGGGG |
| **65** | SP1RACEMS | CGTCTTCGACTTGTGCGGATTTC |
| **66** | SP3RACEMS | CCGTATTCCATAACCTTTACAATCTCGACG |

The underlined bolded fragments of the sequences indicate sites recognized by restriction enzymes attached to oligonucleotides. Positions of the changed nucleotides and introduced mutations are highlighted by small bolded fonts. FAM (fluorescein) is attached to the 5’-end of the oligonucleotide.
